# Single-cell Transcriptomic Profiling Unveils Cardiac Cell-type Specific Response to Maternal Hyperglycemia Underlying the Risk of Congenital Heart Defects

**DOI:** 10.1101/2021.05.28.446177

**Authors:** Sathiyanarayanan Manivannan, Corrin Mansfield, Xinmin Zhang, Karthik. M. Kodigepalli, Uddalak Majumdar, Vidu Garg, Madhumita Basu

## Abstract

Congenital heart disease (CHD) is the most prevalent structural malformations of the heart affecting ∼1% of live births. To date, both damaging genetic variations and adverse environmental exposure such as maternal diabetes have been found to cause CHD. Clinical studies show ∼fivefold higher risk of CHD in the offspring of mothers with pregestational diabetes. Maternal pregestational diabetes affects the gene regulatory networks key to proper cardiac development in the fetus. However, the cell-type specificity of these gene regulatory responses to maternal diabetes and their association with the observed cardiac defects in the fetuses remains unknown. To uncover the transcriptional responses to maternal diabetes in the early embryonic heart, we used an established murine model of pregestational diabetes. In this model, we have previously demonstrated an increased incidence of CHD. Here, we show maternal hyperglycemia (matHG) elicits diverse cellular responses during heart development by single-cell RNA-sequencing in embryonic hearts exposed to control and matHG environment. Through differential gene-expression and pseudotime trajectory analyses of this data, we identified changes in lineage specifying transcription factors, predominantly affecting *Isl1*^+^ second heart field progenitors and *Tnnt2^+^* cardiomyocytes with matHG. Using *in vivo* cell-lineage tracing studies, we confirmed that matHG exposure leads to impaired second heart field-derived cardiomyocyte differentiation. Finally, this work identifies matHG-mediated transcriptional determinants in cardiac cell lineages elevate CHD risk and show perturbations in *Isl1*-dependent gene-regulatory network (*Isl1-GRN*) affect cardiomyocyte differentiation. Functional analysis of this GRN in cardiac progenitor cells will provide further mechanistic insights into matHG-induced severity of CHD associated with diabetic pregnancies.

## Introduction

Congenital heart disease (CHD) is the most common developmental malformation in humans and remains the leading cause of birth-defect related infant mortality. CHD has multifactorial etiology, and the heterogeneous cardiac lesions indicate involvement of both genetic and environmental contributors^1–4^. The genetic basis of CHD has undergone significant investigation, and a disease-causing genetic abnormality is identified in ∼20-30% of all CHD cases ^5–8^. Yet, identification of genetic factors does not inform on the phenotypic variability among CHD patients with identical genetic variants. In addition, studies have identified different environmental stressors such as maternal pre-existing illnesses, viral infections, and therapeutic and nontherapeutic drug exposures^9–12^ that promote CHD. A strong correlation has been identified between maternal pregestational diabetes mellitus (matPGDM) and an increased occurrence of CHD. Estimates from the Center for Disease Control (CDC) reflect that ∼8% of CHD is likely a result of uncontrolled maternal diabetes that is present before and during the first trimester of pregnancy^13, 14^. Therefore, it remains critical to understand the cardiac molecular responses to maternal hyperglycemia (matHG) which compromises CHD-risk loci to increase the incidence of the disease and explain the cardiac lesions in patients harboring identical genetic variants.

Epidemiological studies have suggested that the subtypes of CHD found in infants exposed to matPGDM range from transposition of great arteries, outflow tract (OFT) defects with normally related great arteries, double outlet right ventricle (DORV), cardiac septal defects (ASD, VSD, AVSD) to hypoplastic left heart syndrome^14–16^. Additionally, previous studies from our group and others using animal models of matPGDM have recapitulated human CHD phenotypes and discovered the significance of gene-environment interaction in mouse models contributing to diabetic embryopathy^17, 18^. The prevalence of conotruncal defects in the offspring of matPGDM suggests that hyperglycemic exposure affects cardiac structures that correspond to the second heart field (SHF) in early embryonic development^14, 16^. SHF or pharyngeal mesodermal progenitor cells are situated medial to the primary or first heart field (FHF) and are distinguished by the expression of genes encoding the transcription factors (TFs) Isl1 and Tbx1 and the growth factors Fgf8 and Fgf10^19, 20^. Gene expression, Cre- lineage and retrospective clonal analysis studies in avian models and mouse embryos have demonstrated that SHF cells are multipotent in nature and give rise to the OFT, right ventricle, and atrial cardiomyocytes (CMs) of the heart^19, 21, 22^ with additional contributions to the smooth muscle cells and endocardial/endothelial cells. The specification of SHF progenitor cells and their differentiation to CMs are a tightly regulated process, mediated by a coordinated series of chromatin rearrangements and transcriptional regulation^19, 23^. Although, matHG-mediated transcriptional changes and perturbations of SHF and their derivatives are not completely understood in the context of matPGDM associated CHD. Earlier studies have indicated that direct or indirect effects on the gene-regulatory program driven by TFs (Isl1, Tbx1, Prdm1, Six1, Nkx2.5, Gata4, Mef2c, Hand2), intercellular signaling pathways (Bmp, Fgf, Shh, Wnt, Notch), and chromatin remodeling factors (Smarca4, Smarcd3, Smarcc1) affect SHF deployment and when deleted in mice demonstrate embryonic lethality and higher incidence of CHD^20, 22, 24^. We and others have previously shown that embryonic exposure to matHG is a disruptor of gene-expression of cardiac TFs and epigenetic modifiers, and alters signaling pathways involving Notch, Wnt, Bmp, Tgfβ, Vegf, Shh, Hif1 -pathways in the developing heart^17, 25–27^. The gene-environment interaction studies between matPGDM and *Notch1*, *Nkx2.5*, *Ask1*, *Hif1* haploinsufficiency have also revealed an increased occurrence of VSD, DORV and truncus arteriosus in matHG-exposed embryos compared to embryos exposed to control maternal environment^17, 26–29^. While these studies highlight the need to dissect the molecular mechanism(s) of matHG exposure during cardiac development, the effect of matHG- mediated transcriptional changes in diverse cardiac cell types remain unknown. This impedes precise understanding of the developmental toxicity elicited by matHG in utero.

Here, we used the streptozotocin-induced (STZ) murine model of matPGDM to study the effect of matHG on cellular and molecular changes in developing embryonic hearts. We found direct evidence of altered gene expression upon hyperglycemic exposure *in vivo* using our single-cell RNA-seq (scRNA-seq) analysis in non-diabetic control (CNTRL) and matHG-exposed E9.5 and E11.5 whole hearts. Differential gene expression and gene ontology enrichment analyses have identified HG-mediated changes in *Isl1^+^* multipotent SHF progenitors at E9.5, and in *Tnnt2*^+^ CMs at E9.5 and E11.5. Further, we confirmed our findings from the pseudotime trajectory-based lineage analysis of SHF and CM subpopulations using *in vivo* SHF cell-fate mapping studies. Comparing CNTRL and matHG-exposed *Isl1-Cre^+^; Rosa^mT/mG^* E9.5, E11.5, and E13.5 hearts, we identified that SHF-derived CMs show differentiation defects when exposed to matHG environment. We note that misexpression of cell lineage specifying markers (Isl1, Tbx1, Fgf10, Nkx2-5, Hand2) in scRNA-seq data and flow-sorted E9.5 Isl1+SHF-derived cells indicate that matHG perturbs the Isl1-gene regulatory network (*Isl1-GRN*) to cause CHD. Overall, this study delineates the influence of matHG on CM differentiation defects and scRNA-seq data prioritizes the cellular subtypes that are major contributors to matPGDM-induced CHD.

## Results

### Single cell RNA-sequencing identifies diverse cellular response to matHG in developing E9.5 and E11.5 hearts

To understand the cellular basis of matHG-induced risk of CHD and compare the cardiac cell-type-specific transcriptional responses in CNTRL and matHG-exposed embryonic hearts, we applied *in vivo* 10XscRNA-seq. We used a well-established murine PGDM model that exhibits similar pathogenesis associated with human type 1 diabetes mellitus ^27, 30^. Briefly, wildtype C57BL/6J (*wt*) female mice were treated with STZ to destroy the pancreatic β-cells, resulting in elevated blood glucose (B.G.) levels.

STZ-treated matHG and non-STZ-treated CNTRL *wt* females were bred to adult *wt* males. Maternal B.G. levels in the *wt* females at the time of embryo harvest were measured to confirm the hyperglycemic status (>200 mg/dl) after STZ treatment, following our previously published protocol ^27^. Embryos were collected at E9.5 and E11.5 to examine the effect of matHG on critical stages of cardiac development (**Fig. 1A**). Whole hearts were microdissected and pooled from at least six embryos exposed to CNTRL *wt* (maternal B.G. = 145mg/dl at E9.5 and 196mg/dl at E11.5) and matHG *wt* (maternal B.G. = 312mg/dl at E9.5 and 316mg/dl at E11.5) dams. Cardiac tissues from each developmental stage were dissociated and processed to obtain single-cell libraries using the 10xGenomics Chromium controller and 3‟ polyA-based gene expression analysis kit followed by sequencing (**Fig. 1A**). Recently published R Shiny apps „Natian‟ and „Ryabhatta‟ were used for pre-processing and scRNA-seq data analysis ^31, 32^. We captured single-cell transcriptomic data from 3042 and 4632 cells from CNTRL embryos and 4022 and 3785 cells from matHG-exposed E9.5 and E11.5 embryonic hearts, respectively. To classify the cardiac cell clusters, Seurat-based unsupervised clustering and UMAP (Uniform Manifold Approximation and Projection) dimension reduction was performed after combining four samples and at each stage of development (**Fig. S1A and Fig. S2A, B)**. Cells those pass the quality control (QC) metrics including the number of genes detected in each cell (nFeature-RNA), unique molecular identifiers (UMI or nCount-RNA) and <10% of the reads mapped to mitochondrial genes, were used to analyze differentially expressed genes or DEGs (**Fig. S1A-D and Fig. S2C-F)**. The heatmap and UMAP distribution showed that after combining all four samples, we identified 14 clusters (C0-C13) (**Fig. S1B, C**), annotated based on the top five cell-type-specific marker genes obtained from our data and published scRNA-seq datasets on *wt* embryonic hearts^33, 34^. Cells representing the clusters C3, C6, C7, C11 and C12 and C13 expressed endoderm, ectoderm and blood cell markers and therefore were removed from further analysis (**Fig. S1C, E-G**). We applied similar approach at each stage of development, which identified nine clusters and heatmap showed the top five marker genes per cluster (**Fig. S2G, H**). Clusters which represent the blood, endodermal and unknown population of cells were not included. Following the QC steps, the single-cell transcriptomes from a total of 8503 cells were reduced and classified into six broadly defined cardiac populations and used to compare cell-type-specific DEGs between CNTRL and matHG-exposed E9.5 (1989 and 2304 cells) and E11.5 (2271 and 1939 cells) hearts (**Fig. 1B-D**). These clusters represented *Isl1^+^* and *Tbx1^+^* multipotent progenitors (MP), *Tnnt2^+^ and Actc1^+^* cardiomyocytes (CMs), *Cdh5^+^* and *Emcn^+^* endocardial/endothelial (EC), *Postn^+^* and *Sox9^+^* fibro-mesenchymal (FM), *Wt1^+^* and *Tbx18^+^* epicardial (EP), and *Dlx2^+^ and Dlx5^+^* neural crest (NC) cells. The clustering annotation was performed by finding the gene expression signature of each cluster using marker genes that delineate cell identities as described in published scRNA-seq datasets from *wt* embryonic hearts ^32–34^. The UMAP and dot plots showed cluster-specific expression of marker genes used to classify the cell types (**Fig. S3A, B**).

**Fig. 1.**
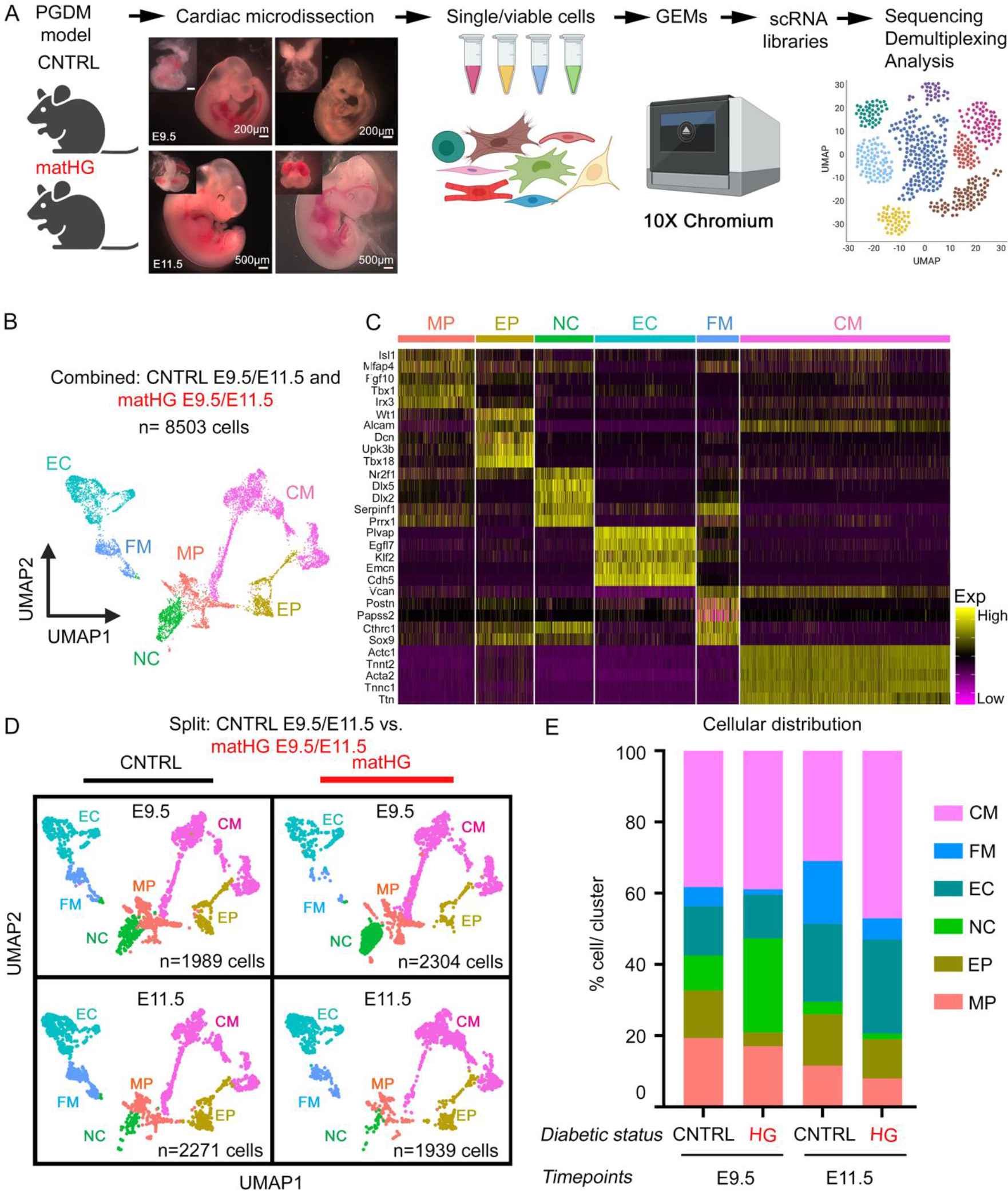
Single-cell transcriptomic sequencing of E9.5 and E11.5 hearts reveal cell type-specific response to maternal hyperglycemic exposure **(A)** Overview of the experimental procedure (made in ©BioRender). E9.5 and E11.5 hearts (n≥6) exposed to CNTRL (B.G <200 mg/dl) and matHG (high B.G >300 mg/dl) dams were microdissected, dissociated to single cells, performed 10X library preparation and sequencing. Dissected tissues are shown in the insets. Scale bars: 200 and 500μm. **(B)** UMAP plot of all cells (n=8503 single cell transcriptomes after combining all four samples) that pass quality control. Cells with similar transcriptional profiles were clustered into six distinct (MP, EP, NC, EC, FM, and CM) cardiac populations. Each dot represents an individual cell and colored according to cluster identities. **(C)** Heatmap showing the expression of top five well-characterized marker genes in each cluster identified from single cell RNA-sequencing. Rows represent cells and ordered by cell cluster identities and hierarchical clustering. Normalized log expression levels are shown in yellow (high expression) and purple (low expression). **(D)** UMAP plots of CNTRL and matHG-exposed cells in each cluster at E9.5 and E11.5 hearts, colored according to the cluster identities. **(E)** Barplot indicating the proportion of cells in each cluster (in percentages) normalized to the total number of cells per sample at E9.5 and E11.5 stages (statistical tests shown in Fig S4A). Cluster identities are indicated in colors. CNTRL, control, HG, hyperglycemia, UMAP, Uniform Manifold Approximation and Projection, MP, multipotent progenitor, EP, epicardial, NC, neural crest, EC, endocardial/endothelial, FM, fibromesenchymal, CM, cardiomyocytes.

Next, comparisons of the cellular proportions from scRNA-seq dataset revealed significant differences between CNTRL and matHG-exposed hearts at E9.5 and E11.5 (**Fig. 1E and Fig. S4A**). We found that CMs constituted the most abundant cell type in E9.5 (CNTRL-38.2% vs. matHG-38.9%) and E11.5 (CNTRL-30.9% vs. matHG-47.1%) and the proportion of CMs was significantly increased in matHG-exposed E11.5 hearts. The MP population in E9.5 (CNTRL-19.3% vs. matHG-17.0%) and E11.5 (11.6% in CNTRL vs. 7.9% in matHG) embryos were both significantly reduced with matHG exposure compared to CNTRL hearts. The FM cells were significantly lower in the E9.5 (CNTRL-5.4% vs. matHG-1.5%) and E11.5 (CNTRL-17.6% vs. matHG-5.9%) hearts subjected to matHG. In contrast, the EC lineage did not change significantly between CNTRL and matHG-exposed E9.5 hearts (CNTRL-13.8% vs. matHG-12.3%) but were significantly increased at E11.5 (CNTRL-21.9% vs. 26.4% in matHG at E11.5). While NC-derived cells in matHG-exposed E9.5 heart (CNTRL-9.8% vs. matHG-26.4%) was significantly greater than CNTRL E9.5 but reduced (CNTRL-3.5% vs. matHG-1.5%) by E11.5 stage (**Fig. 1E and Fig. S4A**). The distribution of EP cells at E9.5 (CNTRL-13.4% vs. matHG-3.9%) and E11.5 (CNTRL-14.4% vs. 11.0% in matHG) between two groups were found to be significantly reduced. These findings from *in vivo* scRNA-seq data identify that matHG exposure was able to induce diverse cellular responses in early stages of cardiac development, predominantly affecting cardiac progenitor cells and their derivatives.

### Maternal hyperglycemia triggers key molecular changes in cardiac progenitors and cardiomyocytes

To gain a better understanding of how matHG exposure was affecting molecular programs involved in heart development, we examined the transcriptional changes in MP and CM clusters from CNTRL and HG-exposed E9.5 and E11.5 scRNA-seq data. The proportion of MP cells show significant differences between CNTRL and matHG-exposed hearts E9.5 and E11.5 hearts while the CMs were significantly altered only at E11.5 (**Fig. 2A and Fig. S4A**). The normalized gene expression of *Isl1^+^*, *Tbx1^+^*, and *Tnnt2^+^* cells in combined MP-CM clusters and triple positive cells indicated the presence of less differentiated CMs (**Fig. 2B**). Bioinformatics analysis revealed 262 DEGs in *Isl1^+^* MP cells at E9.5 with a cutoff value of log_2_FoldChange ≤ -1 or ≥ +1 and P_adj_ ≤0.05 (**Table S1**). Gene Ontology (GO) enrichment analysis with 262 DEGs in E9.5 MP cluster showed significant perturbations in genes associated with biological processes, affecting (i) regionalization, anterior-posterior pattern specification, cell-fate commitment, (ii) cardiomyocyte, mesenchymal and neural crest cell differentiation, and (iii) cardiac ventricle and septum development (**Fig. 2C, Table S2**). The DEGs associated with these processes include Hox-family members, fibroblast growth factors, Forkhead box members, T-box TFs, muscle-specific TFs, and regulators of SHF development (**Table S1**). Therefore, DEG analysis of MP cluster revealed disruption of cardiac progenitor cell commitment and altered CM fate in response to matHG.

**Fig. 2.**
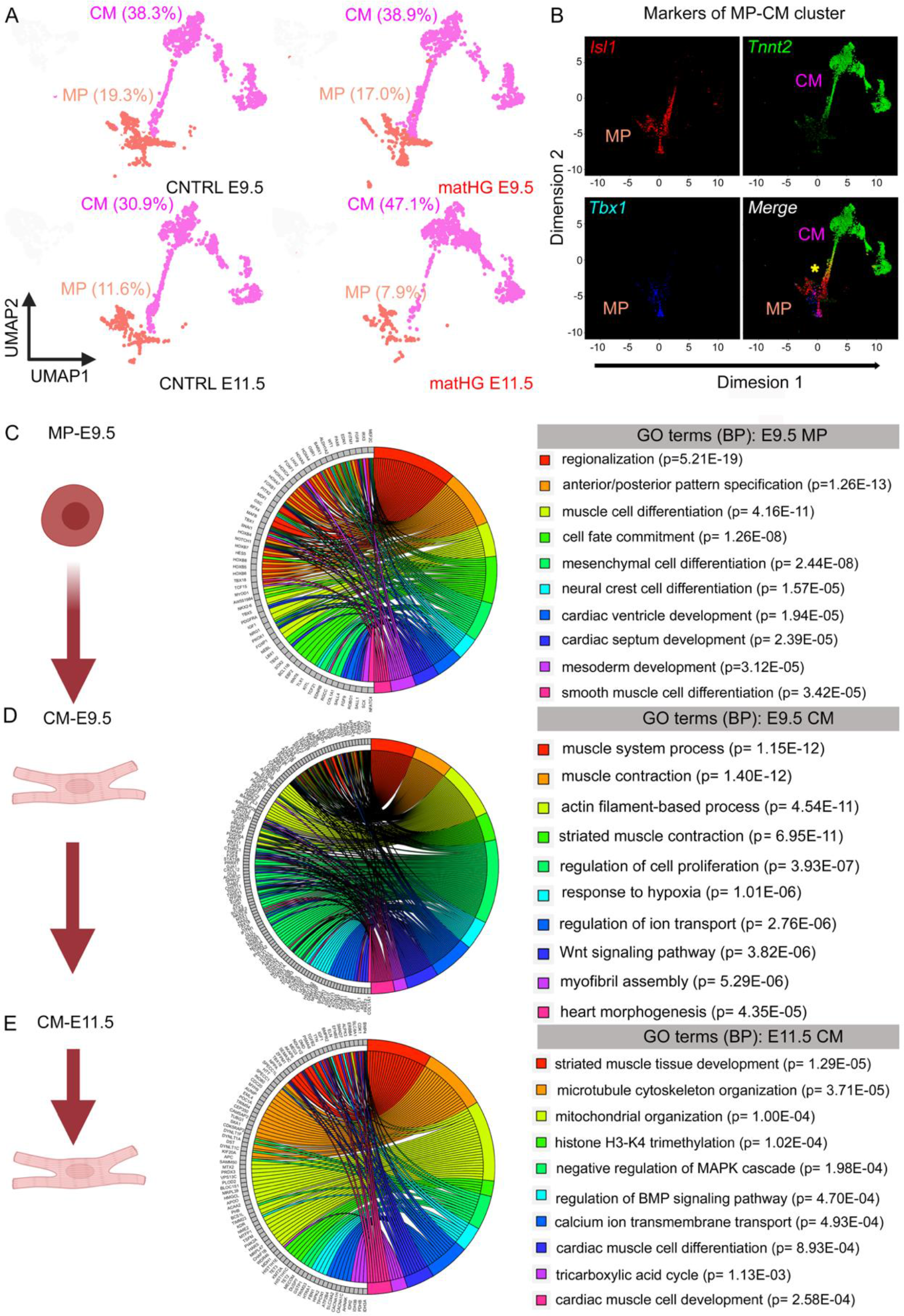
Transcriptomic analysis of matHG-exposed hearts reveals gene expression differences in cardiac progenitor population and in cardiomyocytes **(A)** UMAP plots demonstrate the proportion of cells in MP and CM clusters obtained from CNTRL and matHG-exposed E9.5 and E11.5 hearts. **(B)** ExpressO plot show expression of well-known markers of MP (*Isl1* and *Tbx1*) and CM (*Tnnt2*) clusters. * Indicates presence of *Isl1^+^Tnnt2^+^* cells. **(C-E)** GOplots represent the analysis of the gene ontology (GO) terms enriched among the 262 DEGs in E9.5MP, 357 DEGs in E9.5CM and 326 DEGs in E11.5CM clusters (DEG cutoff: Log2Foldchange ≥ 1 or ≤ -1 and P_adjusted_ ≤ 0.05). The left side of the circle displays the gene, and the right side shows the GO term associated biological processes. The assorted colors represent different GO terms, and the color of each GO terms is annotated, p-values represent enrichment of GO term. CNTRL, control, HG, hyperglycemia, MP, multipotent progenitors, CM, cardiomyocytes, DEG, differentially expressed gene.

Next, the expression analysis was performed in *Tnnt2^+^* CMs between CNTRL and matHG-exposed hearts at E9.5 and E11.5, which revealed 357 and 326 DEGs, respectively (**Tables S1 and S3**). GO enrichment analysis of CNTRL vs. matHG- exposed DEGs in E9.5 and E11.5 CMs showed molecular changes associated with (i) muscle contraction and myofibril assembly, (ii) regulation of ion transport, (iii) cardiac muscle cell differentiation and proliferation, (iv) regulation of Wnt, Bmp, TGF and MAPK signaling pathways, (v) response to hypoxia, (vi) H3K4 trimethylation, and (vii) mitochondrial organization with the regulation of metabolic processes such as tricarboxylic acid cycle (**Fig. 2D, E and Tables S2, S4**). In E9.5 CMs, we found that the DEGs were primarily associated with cell differentiation, voltage-gated calcium channels and potassium channels, regulators of cardiac contractility, transcriptional and chromatin regulators. Likewise, in E11.5 CMs, expression of genes regulating the sarcomere assembly and cardiac muscle function, myocardial TFs, Bmp, TGF and EGF receptor family genes, glucose and mitochondrial metabolism were significantly perturbed with matHG. Together, these results demonstrate that matHG exerts developmental toxic effects in *Isl1^+^* MP and in *Tnnt2^+^* CMs by affecting key biological processes. This data also suggests that deregulated expression of genes important for CM lineage commitment is likely contributing to a spectrum of conotruncal CHD observed in matHG-exposed embryonic hearts.

### Maternal hyperglycemic exposure triggers molecular responses in cardiac progenitor and CM subtypes that underlie the risk of CHD

To determine the effect of matHG on transcriptional differences in MP and CM subpopulations, we performed a sub-clustering analysis on the integrated E9.5 and E11.5 scRNA-seq data (**Fig. 3A**). The UMAP distribution of cells showed two subpopulations of MP and four subpopulations of CMs present in our scRNA-seq data. Unsupervised clustering, dimension reduction and heatmap analysis identified the marker gene expression levels in each subcluster. Based on the expression of top five markers per cluster from previously published scRNA-seq data of *wt* embryonic hearts ^33, 34^, MP and CM clusters were further classified as anterior/posterior SHF and branchiomeric muscle progenitor (BrMP) cells, outflow tract (OFT), atrioventricular canal (AVC), atrial (Atr) and ventricular (Ven)-CM subpopulations (**Fig. 3B**). Expression of marker genes for each CM subtype is shown in dot plots (**Fig. S5A**). The CNTRL and matHG-exposed E9.5 and E11.5 hearts show significant differences in the distribution of MP-CM subpopulations (**Fig. 3C and Fig. S4B**). In presence of matHG exposure, we found a higher proportion of BrMP cells (2.8% in CNTRL vs. 10.8% in matHG) at E9.5, with a decrease in SHF populations both at E9.5 (30.7% in CNTRL vs. 19.6% in matHG) and E11.5 hearts (26.1% in CNTRL vs. 14.1% in matHG). Given the contribution of MP subpopulations during heart development, these cellular differences suggest a potential role of glucose sensitivity in cells from the SHF and increased risk of CHD in the diabetic offspring. This is further reflected in the significant reduction of *Tnnt2^+^* OFT-CMs (18.7% in CNTRL vs. 3.7% in matHG) and AVC-CM (3.3% in CNTRL vs. 0.6% in matHG) at E11.5 with matHG (**Fig. S4B**). We also detected that CNTRL vs. matHG-exposed Atr- (17.0% vs. 14.4%) and Ven-CMs (33.0% vs. 27.6%) were reduced at E9.5, but were significantly higher in the E11.5 hearts (from 16.6% to 26.0% Atr-CM and from 34.2% to 55.1% Ven-CM, respectively) when subjected to matHG environment (**Fig. 3C, D and Fig. S4B**).

**Fig. 3.**
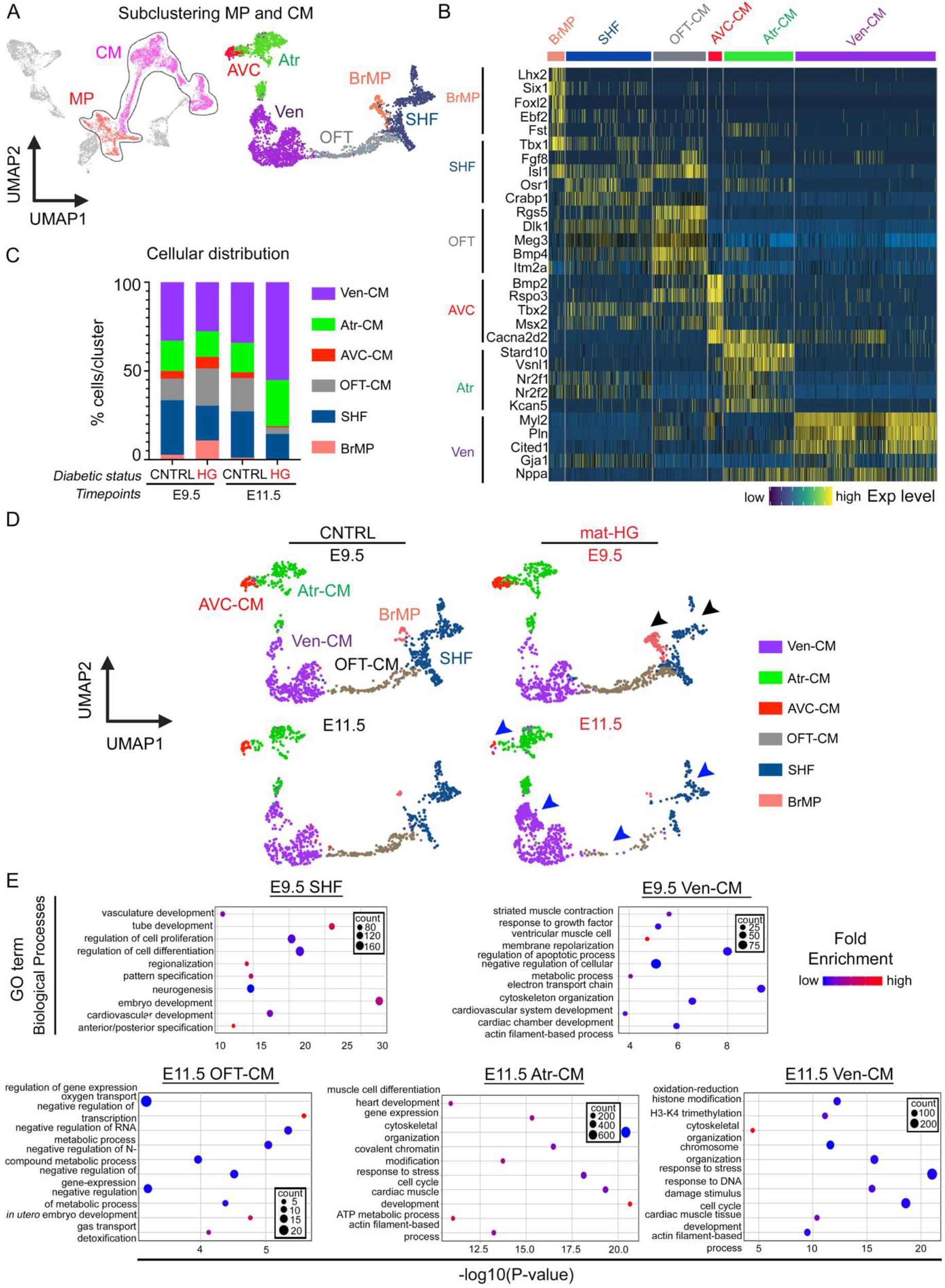
Gene expression profiles of MP-CM cell subpopulations show differential response to matHG **(A)** Combined UMAP plots show subclustering of MP (1194 cells; 14.1%) and CM (3273 cells; 38.5%) clusters (indicated by black dotted line). Cells with similar transcriptional profiles were subclustered into six distinct subpopulations. MP cells were clustered into SHF and BrMP subpopulations and CMs were clustered into OFT-CM, AVC-CM, Atr- CM and Ven-CM subpopulations. Colors denote identity of subclusters. **(B)** Heatmap showing the expression of top five well-characterized marker genes in each subclusters. Rows indicate marker genes and columns denote single cells. **(C)** Barplot indicating the proportion of cells in each subcluster (in percentages) normalized to the total number of cells per sample at E9.5 and E11.5 stages. Cluster identities are indicated in colors (statistical tests shown in Fig S4B). **(D)** UMAP plots of CNTRL and matHG-exposed MP and CM subpopulations at E9.5 and E11.5 hearts, colored according to the cluster identities. Changes in SHF and BrMP cell populations (black arrowheads) in E9.5 and SHF, OFT, Atr and Ven-CMs (blue arrowheads) in E11.5 embryos are indicated. **(E)** Bubble plots represent enriched biological process GO terms in E9.5 SHF and Ven-CM and in E11.5 OFT, Atr and Ven-CM exposed to matHG. The colors of the nodes are illustrated from red to blue in descending order of –log10 (P-value) and fold enrichment. The count represents gene number, also indicated by circle size, whereas the color denotes the up (red) or downregulation (blue) of the specific GO term in the cellular subpopulations. The horizontal (X) axis represents the –log10 (P-value), and the vertical (Y) axis represents the GO terms. CNTRL, control, HG, hyperglycemia, MP, multipotent progenitors, SHF, anterior/posterior second heart field, BrMP, branchiomeric muscle progenitors, OFT, outflow tract, AVC, atrioventricular canal, Atr, atrial, Ven, ventricular, CM, cardiomyocytes, GO, gene ontology.

To characterize matHG-mediated transcriptional changes in the CM subpopulations, we evaluated the expression of marker genes in four CM subpopulations across two developmental timepoints (**Fig. S5B**). The DEG analysis revealed significant downregulation of *Myl2^+^, Gja1^+^, Pln^+^* and *Cited1^+^* cells in Ven-CMs, *Bmp4^+^ and Itm2a^+^* cells in OFT-CMs, *Bmp2^+^, Rspo3^+^* and *Tbx3^+^* cells in AVC-CMs and *Nr2f1^+^* Atr-CMs at E9.5 (**Fig. S5B**). Likewise, in E11.5 hearts, the expression of *Gja1^+^* Ven-CMs, *Bmp4^+^, Rgs5^+^ and Dlk1^+^* OFT-CMs, *Cacna2d2^+^, Tbx3^+^* AVC-CMs and *Kcan5^+^*, *Nr2f1^+^*, *Nr2f2^+^* Atr-CMs were reduced with matHG. The cardiac response to teratogens and concomitant downregulation of TFs important for SHF deployment (*Nkx2-5*, *Mef2c*, *Gata4* and *Tbx5*) ^35–38^ and CM structural genes suggest why matHG- driven transcriptional changes are frequently associated with CHD indicated by studies in humans and animal models. This data reveals that elevated glucose levels during pregnancy can affect cardiomyocyte maturation by altering the expression of key CM genes.

Next, GO enrichment analysis was performed using MP-CM DEGs and significantly enriched biological processes (FDR adjusted p-value ≤0.05) were compared between two developmental timepoints (**Fig. 3E, Table S5-S7**). The DEGs in E9.5 SHF-derived cells were enriched in biological processes related to cell proliferation/differentiation; anterior-posterior pattern specification, as described earlier. While the DEGs in Ven-CM displayed changes in chamber development, muscle cell membrane polarization, muscle contraction. In contrast, OFT, Atr and Ven-CMs at E11.5 hearts showed significant differences in gene expression related to stress response, muscle cell differentiation, muscle contraction, ATP-dependent metabolic processes, chromatin modification and cell cycle genes (**Fig. 3E, Table S5-S7**). Therefore, DEG and GO enrichment analysis between CNTRL and matHG-exposed MP-CM subtypes show that matHG disrupts the critical stages of heart development by affecting genes related to CM differentiation.

### Pseudotime trajectory analysis discovers the effect of matHG exposure on MP- CM lineage

To investigate the impact of matHG on transcriptional differences observed in MP-CM cell lineages, we performed pseudotime ordering of CNTRL and matHG- exposed E9.5 and E11.5 cells using both Monocle (version 2.0) and Slingshot (version 1.8.0). The pseudotime analysis revealed five distinct cell states (States 1-5) (**Fig. 4A**). Overlaying the cluster identities to the combined pseudotime trajectory revealed the distribution of SHF, BrMP, OFT, AVC, Atr and Ven-CMs in each state. State 1 was comprised of 70.8% SHF, 10.9% BrMP, 18.0% OFT-CM, 0.1% Atr-CM, 0.2% Ven-CM.

**Figure 4.**
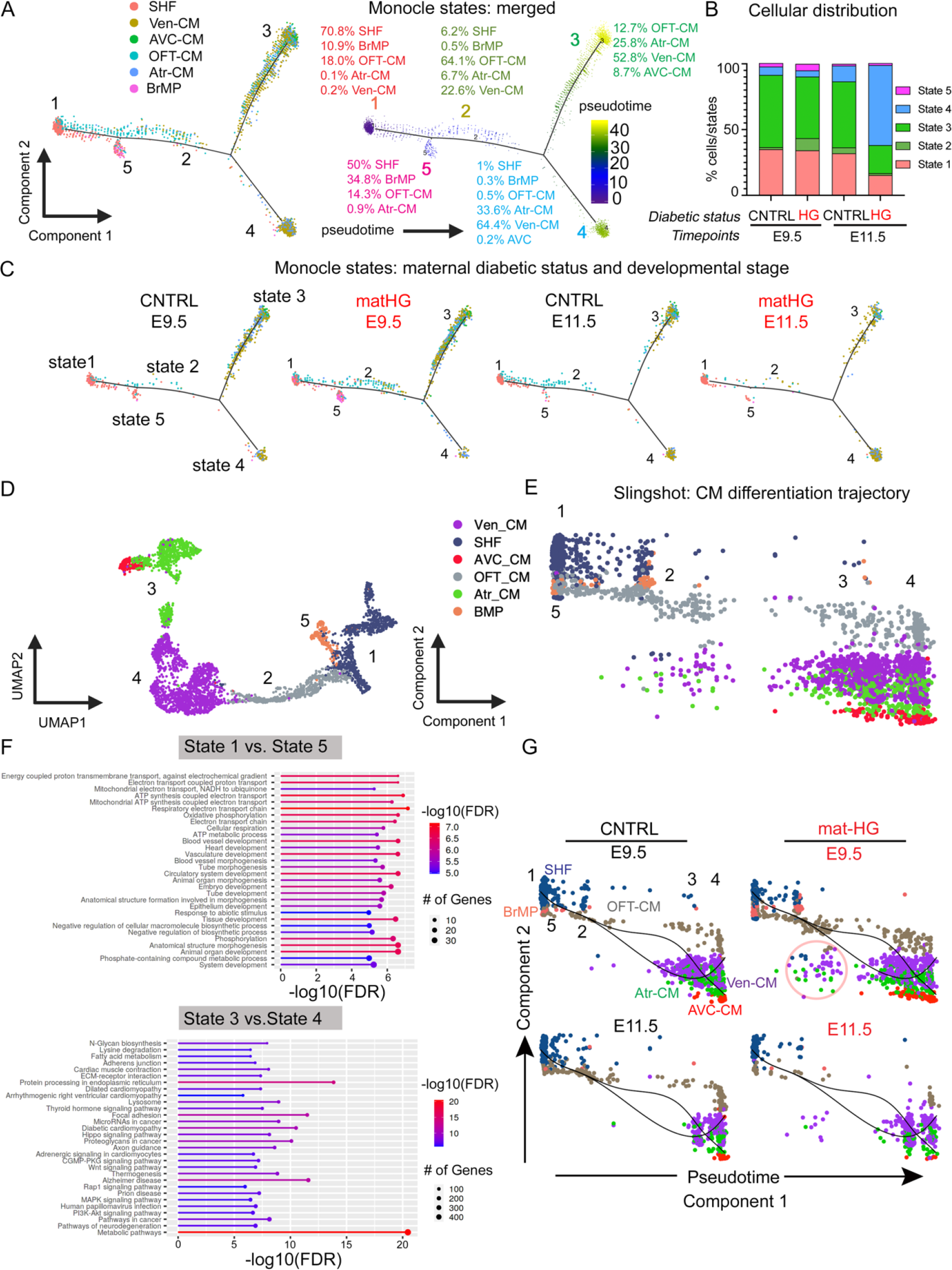
Single-cell pseudotime analysis reveals CM differentiation trajectory from second heart field progenitor cells in response to matHG exposure **(A)** Pseudotime analysis of single cells from combining CNTRL and matHG-exposed E9.5 and E11.5 MP and CM subpopulations (SHF, BrMP, OFT, AVC, Atr and Ven-CM) using Monocle 2. Cells on the tree are colored according to cluster identities, state, and expression levels in pseudotime. The top 500 genes with the highest variability in expression were used to construct the pseudotime tree. The arrangement of cells shows the cells on the left side of the tree (dark blue) are less differentiated (progenitor like) than those on the right side (bright yellow) are more differentiated (CM subpopulations). Overlaying the cluster information shows that cells in states 1 and 5 correspond to the SHF progenitor cells and states 2, 3 and 4 correspond to less differentiated and more differentiated CMs, shown by percentages in each state. **(B)** Barplot indicating the proportion of cells in MP-CM subpopulations (in percentages) across five pseudotime states (1-5) for CNTRL and HG-exposed E9.5 and E11.5 embryos (statistical tests shown in Fig S4C). **(C)** Pseudotime trajectories shown at two developmental stages exposed to CNTRL and matHG conditions. **(D, E)** Slingshot-based pseudotime trajectories calculated from UMAP embedding illustrate the trajectories of MP-CM subpopulations in CNTRL and matHG exposed E9.5 and E11.5 hearts (combined). **(F)** Gene Ontology enrichment analysis of differentially expressed genes between States 1 and 5 (progenitor like) and States 3 and 4 (differentiated CM) were performed using ShinyGO v0.741. (G) Slingshot-based pseudotime trajectories of MP-CM subpopulations in CNTRL and matHG exposed E9.5 and E11.5 hearts (split). Each dot represents an individual cell and is colored according to subcluster identities across the pseudotime. The black line in **(G)** indicates direction of the trajectory. The red circle in matHG-exposed E9.5 trajectory indicates less differentiated CMs at State 2. Overlaying the state and cluster identities show differences between progenitor cells and differentiated CM lineages. MP, multipotent progenitor, CM, cardiomyocytes, CNTRL, control, HG, hyperglycemia, BrMP, branchiomeric muscle progenitors, OFT, outflow tract, AVC, atrioventricular canal, Atr, atrial, Ven, ventricular.

State 5 constitutes 50% SHF, 34.8% BrMP, 14.3% OFT-CM, 0.9% Atr-CM, suggesting that these two states are more progenitor-like. However, we found State 2 contains 6.2% SHF, 0.5% BrMP, 64.1% OFT-CM, 6.7% Atr-CM and 22.6% Ven-CM suggesting an intermediate trajectory for differentiating CMs. In contrast, the State 3 branch was only composed of 12.7% OFT-CM, 25.8% Atr-CM, 52.8% Ven-CM, 8.7% AVC-CM with no expression of SHF and BrMP cells. Finally State 4 constitutes 1% SHF, 0.3% BrMP, 0.5% OFT-CM, 33.6% Atr-CM, 64.4% Ven-CM, 0.2% AVC. The distribution of cells in States 3 and 4 suggested the presence of more differentiated CMs (**Fig. 4A**). The differences in the percentage of cells/state were found to be statistically significant between two developmental time points when exposed to matHG environment (**Fig. 4B and Fig. S4C**). At E9.5, we found significant differences in States 2, 3 and 5 with matHG, while changes in cellular distribution across the trajectory were shown to be significant in States 1-4 at E11.5 (**Fig. 4B, C and Fig. S4C**). Next, Slingshot-based pseudotime trajectories were estimated from UMAP embedding to smoothen the trajectories between MP and CM subpopulations in CNTRL and matHG exposed E9.5 and E11.5 hearts (**Fig. 4D, E**). Like Monocle2, this trajectory analysis revealed progenitor-like States 1 and 5 (far left in the pseudotime), intermediate State 2 with less differentiated CMs and more differentiated CMs at States 3 and 4 (far right in the pseudotime), superimposed with cluster identities. GO enrichment analysis between States 1 and 5 and States 3 and 4 identified DEGs associated with heart development, tube morphogenesis, metabolic processes, and cardiac muscle contraction, distinctly separating the progenitor and CM lineages (**Fig. 4F**). The principal component analysis (PCA) plots for CNTRL and matHG-exposed E9.5 and E11.5 were shown according to their true pseudotime, where trajectories were inferred by Slingshot (**Fig. 4G**). We highlighted the differences noted in intermediate State 2 at E9.5 (indicated by a red circle), suggesting that there is an enriched population of less differentiated CMs in the setting of matHG exposure (**Fig. 5G**). Trajectory-based DEG analysis of States 1-5 was performed at two points in time (**Tables S8, S9**). The GO enrichment analysis of DEGs revealed top 20 GO terms (biological processes) for each pseudotime state. We observed that matHG affected genes essential for cardiac development, tube morphogenesis, cell differentiation, muscle structure development and changes in oxidative phosphorylation and ATP dependent metabolic processes (**Fig. S6A-E, and Fig. S7A-E**). These findings from pseudotime trajectory-based molecular analysis reveal that matHG exposure sensitizes MP-CM lineage during CM differentiation.

**Fig. 5.**
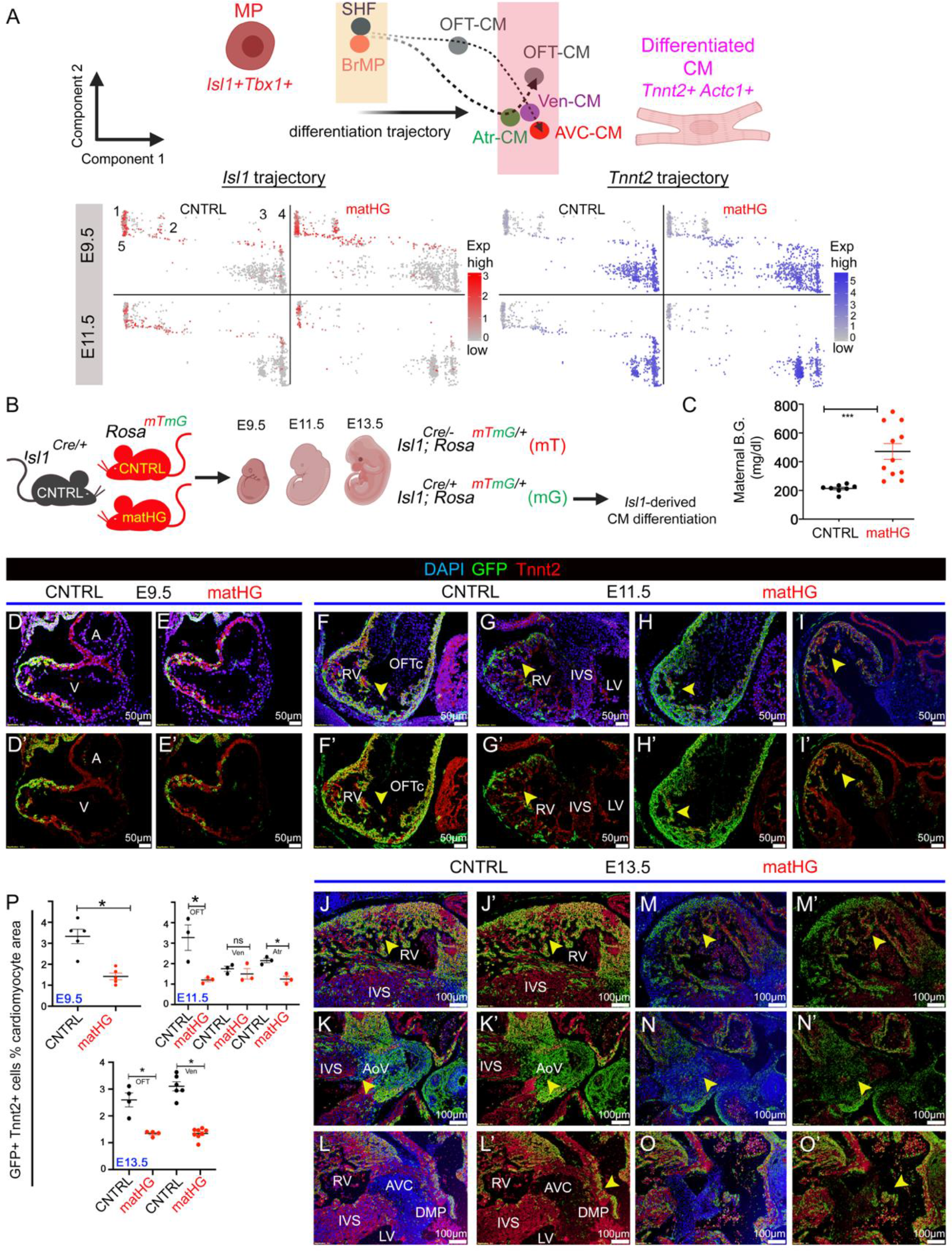
Exposure to matHG impedes SHF-derived cardiomyocyte differentiation *in vivo* **(A)** Schematic showing the trajectory of *Isl1^+^Tbx1^+^* multipotent SHF cells to *Tnnt2^+^Actc1^+^* differentiated CMs. **(B)** Feature plots show the expression profile of SHF- marker *Isl1* and CM-marker *Tnnt2* in CNTRL vs. matHG-exposed E9.5 and E11.5 hearts across five states (state 1-5) in Slingshot-based pseudotime trajectories. Each dot represents single cells and color intensities (red for *Isl1* and blue for *Tnnt2*) display gene expression levels. **(B)** Schematic outlines the experimental strategy for SHF- derived CM lineage analysis in response to matHG. *Isl1-Cre^+/-^* males bred to CNTRL, and STZ-treated matHG *Rosa^mT/mG^* reporter females and embryos collected at E9.5, E11.5 and E13.5 stages for cell fate analysis. Cre^-^ littermates were used as internal control. **(C)** Comparison of maternal B.G. levels (mg/dl) in CNTRL (n=8 litters) vs. STZ- treated *Rosa^mT/mG^* (n=11 litters) dams at the time of embryo harvest. **(D-O)** Lineage-tracing studies in CNTRL and matHG-exposed *Isl1Cre^+^; Rosa^mTmG/+^* embryos show the Isl1-derived GFP^+^Tnnt2^+^ CMs (yellow cells, shown in arrowheads) in E9.5 **(D-E)**, E11.5 **(F-I)** and E13.5 **(J-O)** hearts. **(D’-O’)** Co-immunofluorescence images of GFP^+^ (Isl1- derived, in green) and Tnnt2^+^ (CMs, in red) cells in E9.5, E11.5 and E13.5 embryonic hearts exposed to CNTRL and matHG-environment. Nuclei stained with DAPI shown in blue. **(P)** Quantification of *Isl1Cre^+^; Rosa^mTmG/+^* genetically labeled GFP^+^ cells co-stained with Tnnt2^+^ CMs at three developmental stages (n≥3 per timepoint) exposed to CNTRL and matHG environment. SHF, second heart field, CM, cardiomyocytes, CNTRL, control, HG, hyperglycemia, A, atria; V, ventricle; LV, left ventricle; RV, right ventricle, OFT, outflow tract, IVS, interventricular septum, AoV, aortic valve, AVC, atrioventricular cushion, DMP, dorsal mesenchymal protrusion. Data presented as mean ± SEM; Statistical comparisons made between CNTRL and matHG groups by unpaired Student‟s *t*-test using GraphPad Prism 8. * and *** indicates p-values < 0.05 and <0.001 respectively. Scale bars: 50μm **(D-I)** and 100μm **(J-O)**. Schematic diagrams in **(A, B)** are made in ©BioRender.

### SHF cell-fate mapping analysis demonstrates impaired cardiomyocytes differentiation *in vivo* under matHG

The pseudotime trajectory based DEG analysis in E9.5 and E11.5 hearts revealed the differences in *Isl1^+^Tbx1^+^* MP (progenitor like, States 1 and 5) and *Tnnt2^+^Actc1^+^* CMs (differentiated CMs, States 2-4) (**Fig. 5A**). To examine the role of matHG on the fate of SHF-derived CM differentiation, we first confirmed the *Isl1* and *Tnnt2* gene expression along the pseudotime trajectory using our scRNA-seq data. We found higher number of *Isl1^+^Tnnt2^+^* cells at the less differentiated intermediate State 2 in matHG-exposed E9.5, while the reduction in *Tnnt2^+^* cells was observed in more differentiated CMs at E11.5 State 4 (**Fig. 5A**). This *in silico* prediction suggested that SHF-derived CM differentiation might be affected in response to matHG exposure. To further validate the scRNA-seq prediction and examine if *Isl1*-derived CM differentiation is affected *in vivo*, we used the *Isl1-Cre^+/-^* and *Rosa^mT/mG^* dual reporter mice for cell lineage tracing. CNTRL and STZ-treated matHG *Rosa^mT/mG^* females were bred with *Isl1- Cre^+^* males and E9.5, E11.5, and E13.5 embryos were collected to analyze CM differentiation (**Fig. 5B**). The average maternal B.G. levels between CNTRL (n= 9, 217.7 ± 27.7 mg/dl) and matHG-exposed (n=11, 471.3 ± 182.9 mg/dl) groups were found to be statistically (two-tailed p-value = 0.0007) significant (**Fig. 5C** and **Table S10**). The expression of GFP between CNTRL and matHG-exposed E9.5-E13.5 *Isl1- Cre^+^; Rosa^mTmG/+^* embryos were shown in **Fig. 8A**. This pattern of GFP expression recapitulates the previously described endogenous *Isl1* expression in pharyngeal mesoderm, cardiac OFT, and foregut endoderm at E9.5 and show further extension to the regions of midbrain, forebrain, all cranial ganglia, spinal motor neurons, dorsal root ganglia, and in the posterior hindlimb of E11.5 and E13.5 *Cre^+^* embryos ^39^. Next, we performed co-immunostaining experiments in the E9.5-E13.5 transverse tissue sections with α-GFP (to label Isl1-derived cells) and α-Tnnt2 (to map *Isl1^+^*SHF-derived CMs) exposed to CNTRL and matHG. The cell-fate mapping studies revealed significant reduction in the number of GFP^+^Tnnt2^+^ cells in matHG-exposed E9.5 hearts compared to CNTRL *Cre^+^* embryos (**Fig. 5D-E, P**). The *Isl1*-driven expression of GFP reporter was further examined at later time points (E11.5 and E13.5) and compared between CNTRL and matHG-exposed *Cre^+^* embryos. The Isl1-derived CMs at matHG exposed E11.5 hearts were found to be consistently downregulated in OFT and atria with a trend of downregulation in the RV (**Fig. 5F-I, P**). At E13.5, we found significant downregulation of GFP^+^Tnnt2^+^ cells in both the OFT and RV (**Fig. 5J-O, P**). In addition to CMs, Isl1^+^ SHF cells also give rise to ECs ^40^. Therefore, we checked the effect of matHG on SHF- derived EC cells by plotting *Emcn* expression from scRNAseq data (**Fig. S8B, C)**. The transcript expression was not significantly changed with matHG, however we detected very few *Isl1^+^Emcn^+^* cells *in vivo*. The immunohistochemical analysis of the cardiac sections showed no obvious differences in GFP^+^Emcn^+^ cells marking SHF-derived ECs in matHG-exposed *Is1l-Cre^+^; Rosa^mTmG/+^* E9.5 and E11.5 embryos (**Fig. S8D-K**).

Together, our *in vivo* lineage tracing studies demonstrate that the *Isl1^+^*-SHF-cells were sensitive to the matHG environment and result in impaired CM differentiation.

### Perturbations of *Isl1*-gene regulatory network in response to matHG

To gain insight into HG-mediated changes in the *Isl1*-dependent gene regulatory network (Isl1-GRN) during cardiac development, we reconstructed a protein-protein interaction (PPI) map using the STRINGv11.5 database. We created a list of 34 genes (in mice and humans) encompassing lineage specifying TFs, Fgfs and epigenetic modifiers previously reported ^24^ to function as part of an Isl1-dependent core network for RV and OFT development (**Fig. S9A-D**). The PPI network (enrichment p-value = 1.0E-16) was shown in **Fig. 6A** with Isl1 as a node. We found significant gene expression changes (P_adj_ < 0.05) of the components of Isl1-GRN in E9.5 SHF with matHG exposure, including *Isl1*, *Mef2c, Id2, Pdgfra, Pitx2, Mpped2, Foxp1, Crabp2, Tgfbi, Irx3, Bmpr2, Hes1* (**Fig 6B**). Previous evidence supports that this cardiac TFs individually or in combination with epigenetic modulators regulate SHF development and affect the expansion of CMs ^24, 34, 41, 42^. To test the effect of matHG on Isl1-GRN expression *in vivo*, we focused on two specific candidates, Hand2 and Nkx2-5. Their expression was examined in CNTRL vs. matHG-exposed *Isl1-Cre^+^; Rosa^mTmG/+^* E9.5 hearts. Immunostaining with Hand2 and GFP revealed downregulation of Hand2 expression in matHG-exposed E9.5 OFT (**Fig 6C-H**). While Nkx2-5^+^GFP^+^ expression was found to be downregulated in the OFT and RV, suggesting that matHG could result in abnormal specification of CMs derived from SHF lineage (**Fig 6I-N**). Recent scRNA-seq studies in *Hand2*-null embryos have shown failure of OFT-CM specification, whereas RV myocardium was shown to be specified but failed to properly differentiate and migrate ^34^. Earlier studies have also demonstrated that *Nkx2-5* is required for SHF proliferation through suppression of Bmp2/Smad1 signaling and negatively regulates *Isl1* expression ^42, 43^. The immunohistochemical staining for Mef2c^+^ and Tbx1^+^ cells in matHG-exposed E9.5 *wt* and *Is1l-Cre^+^; Rosa^mTmG/+^* embryonic hearts showed that while Mef2c protein expression was downregulated, Tbx1 expression was greater in the distal OFT in response to matHG (**Fig. S10A-M**). The expression of these cardiac progenitor markers were also measured in FACS-sorted GFP^+^ and GFP^-^ cells from *Is1l-Cre^+^; Rosa^mTmG/+^* E9.5 and E11.5 hearts by qRT-PCR (**Fig. S11A-D**). Significant upregulation of GFP expression (∼83.6-136.9 fold) in CNTRL and matHG-exposed *Is1l-Cre^+^; Rosa^mTmG/+^* embryonic hearts compared to GFP^-^ population confirmed the isolation of two cell populations by FACS (**Fig. S11E**). The qRT-PCR analysis in matHG-exposed E9.5 Isl1- derived GFP^+^ cells revealed significant downregulation of SHF and CM markers including *Isl1, Tbx1, Mef2c, Fgf10, Hand2*, *Nkx2-5, Tbx20, Myl2, Cited2*. While comparisons between CNTRL and matHG-exposed E11.5 GFP^+^ cells demonstrated consistent downregulation of *Tbx1, Mef2c, Hand2*, *Nkx2-5, Myl2,* and *Cited2* and significant upregulation in *Isl1, Fgf10* and *Tbx20* expression at transcript level. Therefore, the altered expression of lineage specifying markers in *Isl1*-derived cells further suggest a potential mechanism for CM differentiation defects contributing to increased risk of CHD in the matHG-exposed embryos. Together, based on our findings, we proposed a “maternal HG-induced CHD model”, in which matHG perturbs Isl1-GRN in the SHF progenitors, leading to impaired CM differentiation and increasing the risk of CHD (**Fig 6O**).

**Fig. 6.**
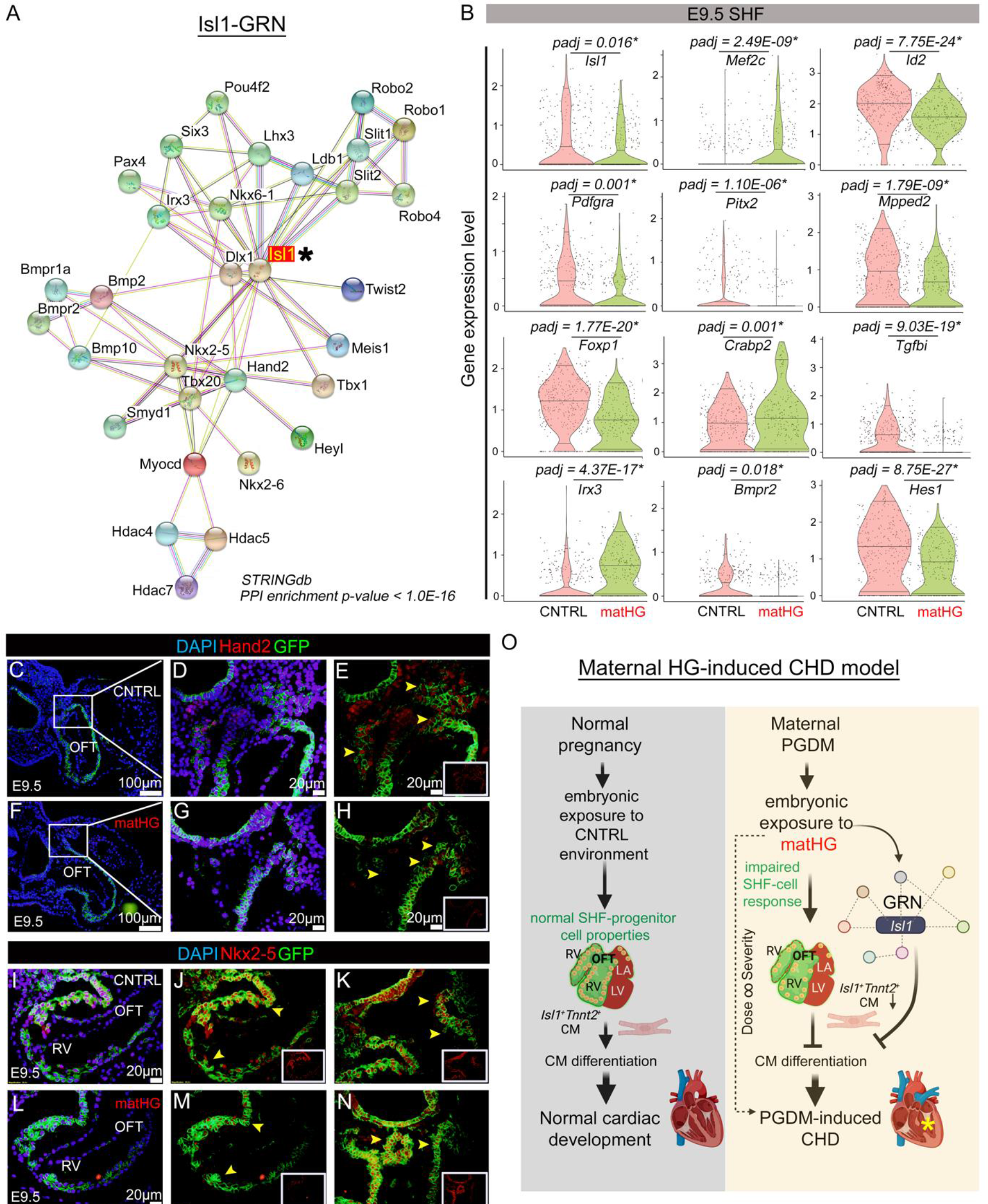
Perturbations in the Isl1-gene regulatory network are intrinsic to matHG- induced CHD **(A)** *Isl1*-dependent gene-regulatory network (Isl1-GRN) created using Isl1-interacting proteins retrieved from published datasets and STRING database representing experimentally established protein-protein interactions (PPI enrichment p-value <1.0E- 16). * Indicates Isl1 as the node. **(B)** Violin plots show normalized gene expression levels of the Isl1-GRN components in CNTRL vs. matHG-exposed SHF cells at E9.5. * Indicates P_adjusted_ (padj) value ≤ 0.05. **(C-N)** Co-immunofluorescence staining shows Nkx2-5 or Hand2 (red, shown in insets) protein expression overlaying with GFP expression driven by Isl1 in E9.5 hearts (n=2) subjected to CNTRL and matHG conditions. The GFP^+^Hand2^+^ cells in E9.5 OFT in CNTRL and matHG-exposed embryos shown in **E, H** (yellow arrowheads). While GFP^+^Nkx2-5^+^ cells were shown in E9.5 OFT and RV-CMs exposed to CNTRL and matHG environment (**J, K, M, N,** yellow arrowheads). Nuclei stained with DAPI shown in blue. **(O)** Proposed model of matHG induced CHD model show impairments in CM differentiation from second myocardial lineage by perturbing genes in Isl1-GRN (made in ©BioRender). Asterisk indicates ventricular septal defects present in matHG-exposed embryos. CNTRL, control, HG, hyperglycemia, OFT, outflow tract, RV, right ventricle, CM, cardiomyocytes, CHD, congenital heart defects. Scale bars: 100μm **(C, F)** and 20μm **(D, E, G, H, and I-N)**.

## Discussion

In this study we report two key findings. First, we have segregated the diverse cellular responses to matHG in the developing mouse hearts using *in vivo* single-cell transcriptomics approach. Second, we extended the computational reconstruction of CM differentiation trajectories to demonstrate that the matHG environment during the early stages of heart development alters the transcriptional network in *Isl1^+^* SHF cells and CM subpopulations. Using cell-fate mapping studies in the matHG background, we were able to confirm that the *Isl1^+^* SHF-progenitor population is susceptible to matHG exposure, and this leads to impaired CM differentiation. In summary, the molecular interactions of Isl1-GRN inferred from single-cell expression data suggest its regulatory role in matPGDM induced CHD.

CHD-causing genes are well studied in murine heart development. Studies have shown that complete deletion of causative genes results in embryonic lethality while heterozygous mice are often unaffected ^44, 45^. Recently, disease modeling of *GATA4, TBX5* pathogenic variants in human iPSC-derived cardiomyocytes and iPSCs-derived endothelial cells from patients with bicuspid aortic valve and calcific aortic valve disease with *NOTCH1* haploinsufficiency have predicted that CHD-linked gene regulatory network is sensitive to the gene dosage ^46–48^. These studies strongly suggest that cell type and gene dosage is critical for normal cardiac development. Like gene dosage, impact of the maternal hyperglycemic status is of high clinical significance. The variability in the adverse matHG environment in presence or absence of genetic variants could similarly impact the disease penetrance and contribute to phenotypic severity observed in CHD patients. The clinical manifestations of matPGDM-induced CHD are variable. Among them, infants born with VSD, conotruncal malformations and heterotaxy are significantly higher in mothers with PGDM ^14, 49^. However, these studies fail to differentiate between the contribution of the genetic variants and that of the maternal environment linked to CHD. The primary teratogen in all human diabetic pregnancies is hyperglycemia, characterized by elevated levels of blood glucose. Additional studies using ex vivo and avian models have confirmed that matHG is the primary teratogen in inducing cardiac defects ^49, 50^. With the help of the experimental model of matPGDM, we and others have previously established a gene-environment interaction model and discovered the increased rate of matHG-induced CHD in presence of haploinsufficient cardiac genes ^11, 26, 27^. More mechanistic studies are needed (i) to identify HG-sensitive cardiac regulatory genes and signaling pathways, (ii) to evaluate their role in FHF, SHF, endocardial/endothelial and cardiac NC cell lineages and (iii) to describe precise mechanism(s) of gene-environment interaction in the setting of matHG. Prior *in vivo* studies with *wt* embryonic hearts led us to compare the breadth of cardiac phenotypes that occur under the direct influence of matHG environment ^27^. Although, future studies are needed to carefully dissect the role of fetoplacental concentration gradient necessary for glucose transfer during the early stages of embryonic development. Better insights into matHG and fetal B.G. levels will provide a robust quantitative measure in defining the cardiac outcome.

The heterogeneity of cardiac lesions observed in the offspring of diabetic mothers suggest a pleotropic effect of matHG on diverse cellular subpopulations. Seminal studies in humans and mice have profiled the single-cell transcriptomes during normal cardiovascular development at an unprecedented resolution to reveal progenitor cell specification and their fate choice ^51–54^. However, the toxicity of matHG (a teratogen) on multiple cardiac cell lineages remain unclear. Using *in vivo* scRNA-seq technology, we first uncovered transcriptional differences in CNTRL vs. matHG-exposed E9.5 and E11.5 hearts. These two points in time were chosen to understand key molecular differences underlying matHG-induced CHD described earlier. After quality control assessment, the unsupervised clustering analysis revealed MP, CM, EC, FM, EP and NC cells in E9.5 and E11.5 hearts based on marker gene expression levels. The differences in cellular distribution between CNTRL and matHG-exposed embryos further confirmed the pleotropic effect of HG. One limitation of this scRNA-sequencing is that we pooled ∼6 embryonic hearts exposed to high-matHG (>300mg/dl) and ∼6 from CNTRL (<200 mg/dl) embryonic hearts for comparative analysis. Previous studies by us and other groups have described that the incidence of matHG-induced CHD (in *wt* embryos) varies from ∼10-50%, so by pooling embryos, we probably underestimated the changes in differential expression from genes with subtle effects.

Based on the manifestation of CHD in the matHG-exposed embryos, we found that a population of *Isl1^+^* cells which marks the SHF show significant gene expression changes upon matHG exposure. The SHF gives rise to the OFT, RV and parts of the Atr-CMs, where the expression of *Isl1* is regulated as the cells adopt a differentiated phenotype ^55^. In human studies, loss-of-function variants in *ISL1* were shown to contribute to CHD and dilated cardiomyopathy alone or in synergy with *MEF2C*, *TBX20* or *GATA4* ^56^. Mutations in this gene were also found to be associated with maturity-onset diabetes of the young and type 2 diabetes patients ^56, 57^. In this study, we demonstrated that the *Isl1^+^* MP and *Tnnt2^+^* CMs subjected to matHG have the most significant differences in gene expression among other cell types identified in scRNA-seq. Therefore, we chose to perform subclustering and pseudotemporal ordering of the MP-CM clusters to identify the role of matHG on CM subpopulations. This data identified gene expression differences in *Six1^+^ Fst^+^* BrMP, *Isl1^+^ Tbx1^+^Osr1^+^* SHF and *Bmp4^+^Rgs5^+^* OFT, *Bmp2^+^Rspo3^+^* AVC, *Myl2^+^Pln^+^* Ven and *Nr2f1^+^Kcna5^+^* Atr-CM subpopulations in matHG-exposed hearts compared to CNTRL. Notably, matHG- induced higher expression of *Isl1* in myogenic progenitors of branchial arches and a reduction in anterior heart field cells suggest that dysregulation of these precursors cells and their derivatives lead to CHD. Monocle and Slingshot based trajectory analysis of scRNA-seq data enabled us to identify progenitor-like States 1 and 5, intermediate and less differentiated CMs at State 2 and more differentiated CMs at States 3 and 4. State-specific DEG and GO enrichment analysis identified several signaling pathways altered in response to matHG exposure include PPAR, BMP, focal adhesion-PI3K-Akt-mTOR, adipogenesis, embryonic stem cell pluripotency pathways, calcium signaling, myometrial relaxation and contraction pathways, HIF1α and p53 signaling, oxidative phosphorylation, pyruvate metabolism, glycolysis/gluconeogenesis. This study provide a list of novel candidate pathways that needs to be evaluated for potential gene-environment interaction in cardiac progenitor cells, and facilitate our understanding of matHG-mediated CHD. To investigate if the SHF-derived CMs are more HG-sensitive, and that CM differentiation is affected, we performed *in vivo* genetic cell fate mapping studies. This data revealed a reduction in *Isl1-*derived GFP^+^Tnnt2^+^ cells in E9.5, E11.5, and E13.5 *Isl1-Cre^+^; Rosa^mt/mG^* embryonic hearts when exposed to matHG. In our previous studies, we have reported CM proliferation defects in E13.5 embryos when subjected to matHG exposure^27^. Future studies are required to address if cardiac proliferation defect precedes the delayed differentiation of CMs using *in vivo* lineage tracing model.

The scRNA-seq data also estimated the differences in gene expression in other cell-types previously associated with CHD. We summarized the DEG and GO enrichment analysis in EC, FM, EP, and NC clusters (**Tables S1-S4**). Our data showed significant expression changes in the genes required for extracellular matrix (ECM) organization, angiogenesis, and tube formation, OFT morphogenesis and heart valve development (**Fig. S12A-I and Fig. S13A-G**). Here, EC-FM genes including *Notch1, Klf4, TGFβ, Snai2, Sox9, Vcan, Twist2, Pdlim3, Gata6, Bmp4, Msx1, Pax3* were shown to be downregulated with matHG. These genes have previously been implicated with endothelial to mesenchymal transition (EndoMT), mesenchymal cell differentiation, and migration of cardiac NC cells ^17, 27, 30^. Significant reduction of FM cells in matHG-exposed E9.5 and E11.5 hearts (**Fig. S4A**) and downregulation of EndoMT genes such as *Vcan Sox9, Postn,* and Bmp/Tgfβ signaling were presented (**Fig. S12E-I**). In the *Tbx18^+^* EP cluster, we showed that the expression of *Wt1, Dcn,* and *Upk3b* were significantly downregulated with matHG and Wnt signaling inhibitor, *Sfrp1* was upregulated at E9.5 (**Fig. S13E**). This data suggest migratory defects in proepicardial derived mesothelial cells. By immunohistochemical staining, we also observed that matHG reduced Fn1 protein expression in E13.5 embryos (**Fig. S13F**). Previous studies have demonstrated that EP-EMT is followed by an activation of Fn1 expression necessary for ECM production and cell migration during heart development ^58^. In the NC cluster, we observed significantly higher expression of *Dlx2^+^, Dlx5^+^* cells only at E9.5 hearts, which was reduced by E11.5 (**Fig. S13E-G**). A recent scRNA-seq study using E8.5-E10.5 *Wnt1^Cre^/R26R^Tomato^* mouse embryos has highlighted the branching trajectory of differentiating NC cells, where expression of *Dlx6* is followed by activation of *Msx2*, *Hand2*, and other cardiac markers such as *Hand1* and *Gata6* ^59^. Therefore, future fate-mapping studies in Wnt1 lineage are necessary that would explain the overabundance of NC population observed in matHG-exposed E9.5 hearts. Therefore, it is important to dissect the underlying cell-type-specific molecular responses in matHG-exposed embryonic hearts. These findings will explain how maternal adverse environment interacts with at-risk loci to increase the risk of CHD in patients harboring identical genetic variants.

Future mechanistic studies are required to evaluate the epigenetic basis of Isl1- GRN under matHG exposure and their progenitor-specific role in regulating CM differentiation (**Fig 6O**). Overall, in this study we reported the embryonic response to matHG environment. The scRNA-seq data uncovered cell-type-specific transcriptional responses to matHG contributing to CHD risk, however future work is warranted to test their causality. The DEG and GO enrichment analysis of each cluster and pseudotime states provide a list of candidates genes and pathways to study gene-environment interaction and facilitate the development of new therapeutics to mitigate matPGDM- induced CHD.

## Methods

### Generation of chemically induced maternal pre-gestational diabetes mellitus (matPGDM) murine model

Wildtype (*wt*) C57BL/6J mice were purchased from Jackson Laboratory (Stock Nos: 000664) for this study. A subset of six- to eight-week-old female mice (∼15-18g body weight) were used to chemically induce type 1-like DM. Mice fasted for an hour before treatment and Streptozotocin (STZ, Fisher Scientific; NC0146241), dissolved in 0.01 mol/l citrate buffer (pH 4.5) was intraperitoneally injected at 75 mg/kg/bodyweight for 3 consecutive days, as previously published ^27^. Seven days post-STZ treatment, mice fasted for ∼8 hours during the light cycle, and blood glucose (B.G.) levels were checked using the AlphaTrak veterinary blood glucometer calibrated specifically for rodents from tail vein blood (Abbott Laboratories). Blood glucose was documented before initiating the timed breeding and during embryo collection. Mice with fasting blood glucose ≥ 200 mg/dl (11 mmol/l) were defined as HG following previously published protocol. If the mice did not achieve the B.G. threshold after 7 days, blood glucose levels were re- tested after fourteen days of STZ injection and then initiated the breeding.

### Dissection of mouse embryos exposed to maternal control and hyperglycemic environment

STZ-treated (matHG) and non-STZ-treated (CNTRL) females were timed bred overnight with *wt* C57BL/6J males. Mice were maintained on a 12-hour-light/dark cycle, and in the morning, when a vaginal plug was observed it indicated embryonic day (E) 0.5. Pregnant mothers were monitored regularly and sacrificed for embryo harvest at E9.5, E11.5, and E13.5 timepoints, and this staging was based on previous literature ^60^. All the mice were housed at the animal facility per Nationwide Children‟s Hospital, Abigail Wexner Research Institute‟s Animal Resource Center policies, and the NIH‟s *Guide for the Care and Use of Laboratory Animals* (National Academies Press, 2011). All animal research has been reviewed and approved by an Institutional Animal Care and Use Committee (protocols: AR13-00056 and AR20-00029).

### Dissection of mouse embryonic hearts and workflow of single-cell RNA-sequencing

To prepare scRNA-seq, the entire cardiogenic region was micro-dissected from E9.5 and E11.5 embryos harvested from *wt* STZ-treated high-matHG and untreated CNTRL female mice. Maternal B.G. levels were tested just before harvesting embryos to ensure diabetic status in CNTRL and STZ-treated females. First, embryos were removed from the yolk-sac, dissected in diethylpyrocarbonate-treated 1X ice-cold PBS, and placed in 1XPBS. Dissected cardiac tissue was incubated in 1mg/ml Collagenase II (Worthington; LS004176) and 1X TrypLE™ Select Enzyme (ThermoFisher Scientific, #12563029) for 15 mins (for E9.5) and 25 mins (for E11.5) at 37°C dry bath with occasional stirring every 5 mins for complete dissociation. The Collagenase/TrypLE solution was quenched immediately with complete DMEM media supplemented with 10% FBS and pelleted down at 1000 rpm for 5 mins at 4°C. Cell pellets were dissolved in 0.04% bovine serum albumin, BSA (Fisher Scientific, #BP9703100) made in PBS and filtered through a 40μm cell strainer (BD Falcon, #352340), centrifuged at 1000 rpm for 5 mins at 4°C, and resuspended in 50μl 0.04% BSA/PBS. Cell viabilities (>85-92%) were assessed using the Trypan blue (1450013; Bio-Rad) exclusion method on Countess^TM^ II FL Automated Cell Counter (ThermoFisher; AMQAF1000). Single-cell libraries targeting ∼4000 cell recovery/sample were generated using 10X Genomics Chromium controller according to the manufacturer‟s instructions using Chromium Single Cell 3′ Reagent Kit (v2 chemistry; PN-120237). The cDNA and libraries were generated using the Chromium Single Cell 3′ Library & Gel Bead Kit v2 and Chromium i7 Multiplex Kit (10X, PN-120237, PN-120262) following the manufacturer‟s suggested protocol. The quality and concentrations of cDNA from each sample were measured using High Sensitivity D5000 ScreenTape® on Agilent 2200 TapeStation. After adjusting all four samples (CNTRL vs. matHG-exposed E9.5 and E11.5) to similar concentrations, each single-cell cDNA sample was used for library preparation and quantified using High Sensitivity D1000 ScreenTape®. Single-cell libraries were sequenced on the Illumina Hiseq4000 platform with 2 X 150 bp read length at the Institute for Genomic Medicine in NCH. CNTRL and matHG-exposed E9.5 libraries were pooled and sequenced in the same lane. Similarly, CNTRL and matHG-exposed E11.5 single-cell libraries were pooled together and sequenced in the same lane. Sequencing parameters were selected according to the Chromium Single Cell v2 specifications. All libraries were sequenced to a mean read depth of at least 50,000 reads per cell.

### Single-cell RNA-seq data pre-processing, quality control and unsupervised clustering

We used the Cell Ranger „mkfastq‟ function to demultiplex and convert Illumina „.bcl‟ output into fastq files. We mapped the fastq reads to the mouse genome mm10 (GRCm38.p6) and gene annotation downloaded from 10X Genomics (https://cf.10xgenomics.com/supp/cell-exp/refdata-gex-GRCh38-2020-A.tar.gz) using the Cell Ranger „count‟ function and generated the gene-count matrix output. The Cell Ranger output was used to create a Seurat object using the R Shiny app Natian (available through www.singlecelltranscriptomics.org). Cells of low quality or those representing doublets were excluded from our analyses using the following cutoffs: Number of genes expressed (*nFeature_RNA*) set to 500-7000 and percentage of mitochondrial gene expression relative to total expression of the cell (*percent.mt* ) < or = 10%, we obtained 2823, 3239, 3836 and 3229 cells from CNTRL-E9.5, CNTRL-E11.5, matHG-E9.5, and matHG-E11.5 respectively. This data was processed using Natian to perform normalization and identify genes with the highest variability ^61^. The data was scaled and dimensionality reduction was performed using Natian and clustering of cells was visualized in two dimensions using Uniform Manifold Approximation and Projection (UMAP)^62^. Briefly, we added „DevStage‟ and „Maternal‟ meta information to each Seurat object to define the developmental stage and the diabetes state of the dam, respectively. Next, we combined the data using regularized negative binomial regression („SCTransform‟ function in Seurat) and canonical correlation analysis (CCA) based integration in Natian. We also tried integration without „SCTransform‟ function and using 50 dimensions and CCA to obtain a merged dataset. There was no significant difference in the clustering of cells based on the two approaches. We performed dimensionality reduction and clustering of cells on the integrated data using a very broad clustering parameter (resolution = 0.2).

### Cell type identification and sub-clustering analysis

Marker genes were identified for individual clusters using a minimum percent expression of 50% and log_2_fold change threshold of 0.25 (log_2_FC.threshold). From each cluster, the top 5 markers were selected based on average log_2_FC and used to classify each cluster in Ryabhatta (available at www.singlecelltranscriptomics.org). The clusters were also cross-referenced with known cell-type-specific marker gene expression using publicly available wildtype scRNA-seq datasets at other embryonic time points ^33, 34^. All sub-clustering analyses (on multipotent progenitor cell and cardiomyocyte subpopulations) were processed using Ryabhatta using a similar number of principal components and resolution parameters.

### Differential gene expression, Gene Ontology enrichment, and protein-protein interaction analysis

Cluster-specific differential gene expression analysis was performed between embryonic developmental stage and maternal diabetic status. Within each group, we combined counts obtained for each gene to produce three in silico replicates of gene vs. expression count matrix. These in silico replicates were then analyzed using DESeq2 to identify differentially expressed genes between various conditions ^63^. Owing to the high drop-out rate observed in the 10X drop-seq method, our approach to combine counts reduces noise and brings the data closer to bulk-RNA seq data which are conventionally used in the DESeq2 pipeline ^63, 64^. The Gene Ontology (GO) annotation of the up/down- regulated DEGs was performed using ShinyGO v0.741 software, a web-based graphical gene-set enrichment tool ^65^. The biological process, affected by matHG exposure was represented as fold enrichment, the number of genes in each GO-term with - log10(FDR) values <0.05. Protein-protein Interaction (PPI) networks (mouse and humans) of identified Isl1-interacting proteins (n=34 proteins) were created using STRING v11.5. STRING database (https://string-db.org) is a curated knowledge database of known and predicted protein-protein interactions ^66^. Most of the Isl1-GRN proteins retrieved from published literature ^24^ and demonstrated an established link with each other in the interaction network.

### Pseudo-time trajectory analysis

Cell trajectory analyses were performed on matHG-exposed E9.5 and E11.5 vs. CNTRL E9.5 and E11.5 multipotent progenitor cell and cardiomyocytes subpopulations using the Monocle 2 (http://cole-trapnell-lab.github.io/monocle-release/) in Ryabhatta. The pseudotime data was further used to generate smooth trajectory curves using Slingshot version 1.8.0 packages ^67, 68^. Differentially expressed genes were determined using the *FindAllMarkers* function in the Seurat package implemented in Ryabhatta for temporal ordering of these cardiac cells along the differentiation trajectory in response to CNTRL vs. matHG environment.

### Second heart field cell-lineage tracing in response to maternal hyperglycemic exposure in utero

For *Isl-1*-derived cardiac cell-lineage tracing studies, we purchased *Isl-1Cre^+/-^* male mice and double fluorescent *Rosa^mT/mG^* reporter mice from Jackson Laboratory (Stock Nos: 024242 and 007676). Adult males were bred with 6-8 weeks old STZ-treated and untreated homozygous *Rosa^mT/mG^* female mice. Diabetes was chemically induced in females as described earlier and B.G. levels were tested 14 days post-STZ treatment. Mice with fasting blood glucose ≥ 250 mg/dl were defined as HG in *Rosa^mT/mG^* strain.

Following successful breeding CNTRL and matHG-exposed *Isl-1Cre^+^; Rosa^mTmG/+^* embryos at E9.5, E11.5, and E13.5 were harvested and compared with *Cre^-^* littermate controls. Embryos were fixed at 4% paraformaldehyde for 24 hours and changed to PBS. TdTomato and GFP fluorescence intensities of whole embryos were captured using an Olympus BX51 microscope.

### Immunofluorescence staining

For immunofluorescence (IF) staining, E9.5, E11.5, E13.5 formalin-fixed paraffin-embedded cardiac sections were deparaffinized using xylene and grades of ethanol, followed by antigen retrieval using citrate-based Antigen Unmasking solution (H-3300, Vector laboratories) using standard protocols ^27^. After permeabilization and blocking with 1% BSA in PBS-Triton X-100 for 1 hour, tissue sections were probed overnight at 4°C with primary antibodies including rat α-Endomucin (1:250; Millipore; MAB2624); mouse α-Cardiac Troponin T (1:250; Abcam; ab8295), rabbit α-Periostin (1:250; Abcam; ab14041), rabbit α-Transgelin or SM22-α (1:250; Abcam; ab14106), rabbit α- Fibronectin (1:250; Abcam; ab2413), and rabbit α-GFP (1:250; ab290; Abcam). Following a series of washing, sections were incubated with a donkey α-rat, α-rabbit, and α-mouse secondary antibodies conjugated to Alexa Fluor 594/488 for 1 hour at room temperature in the dark. After washing, sections were counterstained with Vectashield HardSet Antifade Mounting Medium with DAPI (Vector laboratories). The images were visualized using Olympus BX51 and Zeiss AxioImagerA2. All staining experiments were performed at least in triplicate.

### Tissue dissociation and GFP^+^ cell sorting of *Rosa^mT/mG^* embryonic hearts

The E9.5 and E11.5 embryonic hearts exposed to CNTRL and matHG dams were isolated by microdissection and dissociated to single cells by collagenase digestion as previously described. Isolated cells were FACS-sorted into GFP positive and GFP negative populations using BD FACSAria™ at NCH flow cytometry core. Sorted cells were collected into TRIzol (15596018, ThermoFisher Scientific) and frozen at -20°C for RNA isolation.

### RNA purification and quantitative real-time PCR

RNA was extracted from flow-sorted GFP positive and GFP negative populations exposed to CNTRL and matHG using TRIzol Reagent, as described earlier ^27^. RNA was quantified, and 500ng-1μg of total RNA was used for reverse transcription using the SuperScript VILO cDNA Synthesis Kit (11754-050, ThermoFisher Scientific). SYBR Green-based qRT-PCR was performed for SHF (*Isl1, Tbx1, Mef2c, Fgf10, Hand2*), and CM (*Nkx2-5, Tbx20, Myl2, Cited2*) markers using StepOnePlus™ Real-Time PCR System (Applied Biosystems). Mean relative gene expression was calculated after normalizing Ct values to *Gapdh* using the ΔΔCt method and presented as Log_10_(Fold change). Three independent replicates were performed after pooling 3 embryos per group/timepoint. Oligonucleotide sequences of these genes are provided in **Table S11**.

### Statistical analysis

All experiments were performed at least in triplicate and data represented as mean ± SEM unless otherwise mentioned. Two-tailed Student‟s *t*-test, Fisher exact test and chi-square test with Yate‟s correction (for categorical data) were performed to determine statistical significance using GraphPad Prism 8 software package. *, p ≤ 0.05 is statistically significant.

### Data and materials availability

The authors declare that all supporting data are available within the article and its Supplemental Materials. The scRNA sequencing data is available at the NCBI GEO accession number and the R Shiny apps used to create, analyze, and visualize single-cell transcriptomic datasets are publicly available at www.singlecelltranscriptomics.org ^31^. The mouse models are commercially available from Jackson Laboratory, USA. Any other relevant data are available from the lead corresponding author upon reasonable request. Source data are provided with this paper.

## Author contributions

MB and VG conceived the project and designed and interpreted the experimental results with input from SM for scRNA-seq data. MB supervised the experiments with input from VG. MB performed immunohistochemical characterization, generated single-cell libraries, acquired all the data and analyzed the experiments. SM performed bioinformatics analysis of scRNA-seq data, pseudotime trajectory analysis using Monocle and Slingshot and developed the Seurat-based graphical user interface, Natian and Ryabhatta. CM performed the murine studies including STZ injections, embryo collection, genotyping, sectioning and histological imaging and quantification with supervision by MB. XZ performed initial scRNA-seq data processing. MK generated control E11.5 single-cell library. UM assisted in the analysis of histological images. MB wrote the manuscript with input from SM, CM, VG, and all the other authors.

## Competing interests

Authors declare no competing interests.

## Acknowledgments

The authors thank members of the Biomorphology core for histology support and The Steve and Cindy Rasmussen Institute for Genomic Medicine at Nationwide Children‟s Hospital for sequencing and preliminary bioinformatic analysis. The authors are grateful to Dr. Kedryn K. Baskin for helpful comments and review of the manuscript.

## Funding

This work was supported by funding from the American Heart Association and the Children‟s Heart Foundation Career Development Award Grant 18CDA34110330 (M.B), funding from the National Institutes of Health/National Heart Lung and Blood Institute award R01HL144009 (V.G.) and a Postdoctoral Fellowship award T32HL098039 (S.M).

## Supplemental Materials

**Fig. S1.**
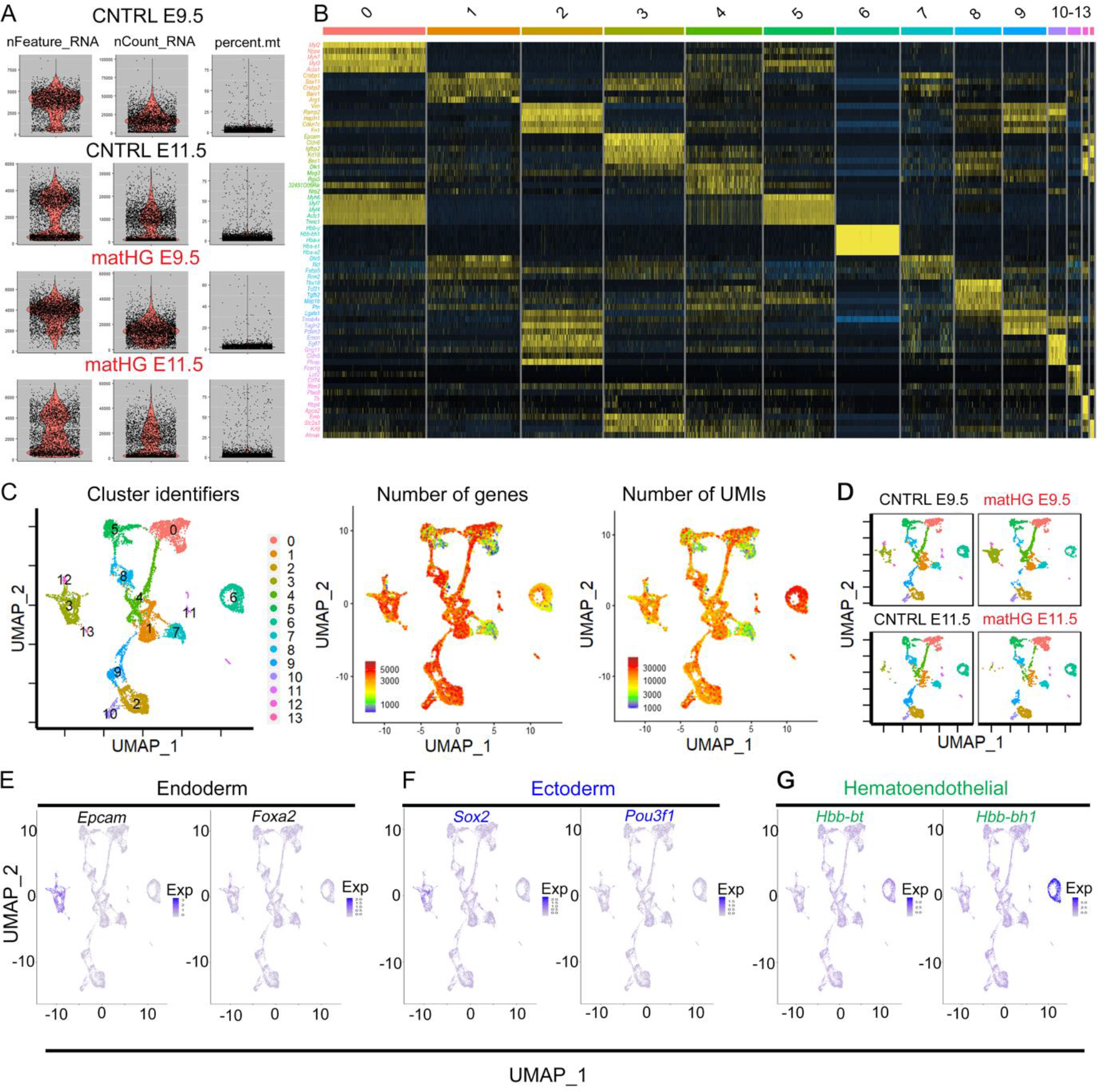
Quality control and normalization of *in vivo* scRNA-seq data **(A)** Quality control metrices used to assess the quality of 10xscRNA-seq libraries from CNTRL and matHG-exposed E9.5 and E11.5 hearts. Violin plots illustrating number of genes detected in each cell (nFeature-RNA), unique molecular identifiers (nCount-RNA) and less than 10% of the reads mapped to mitochondrial genes in each of the four cardiac tissue samples. **(B)** Unsupervised clustering shows a total of 14 clusters (0-13) and the top five marker genes per cluster in the heatmap. Normalized log expression levels are shown in yellow (high expression) and dark blue (low expression). **(C, D)** UMAP plots show the cluster identities and relationship between the number of genes and UMIs in merged datasets and in all four samples. **(E-G)** UMAP plots showing the expression of *Epcam^+^*, *Foxa2^+^* endodermal, *Sox2^+^*, *Pou3f1^+^* ectodermal and *Hbb-bt*^+^*Hbb-bh1^+^* hematoendothelial clusters were discarded from subsequent analysis. CNTRL, control, HG, hyperglycemia, UMAP, Uniform manifold approximation and projection, UMI, unique molecular identifiers.

**Fig. S2.**
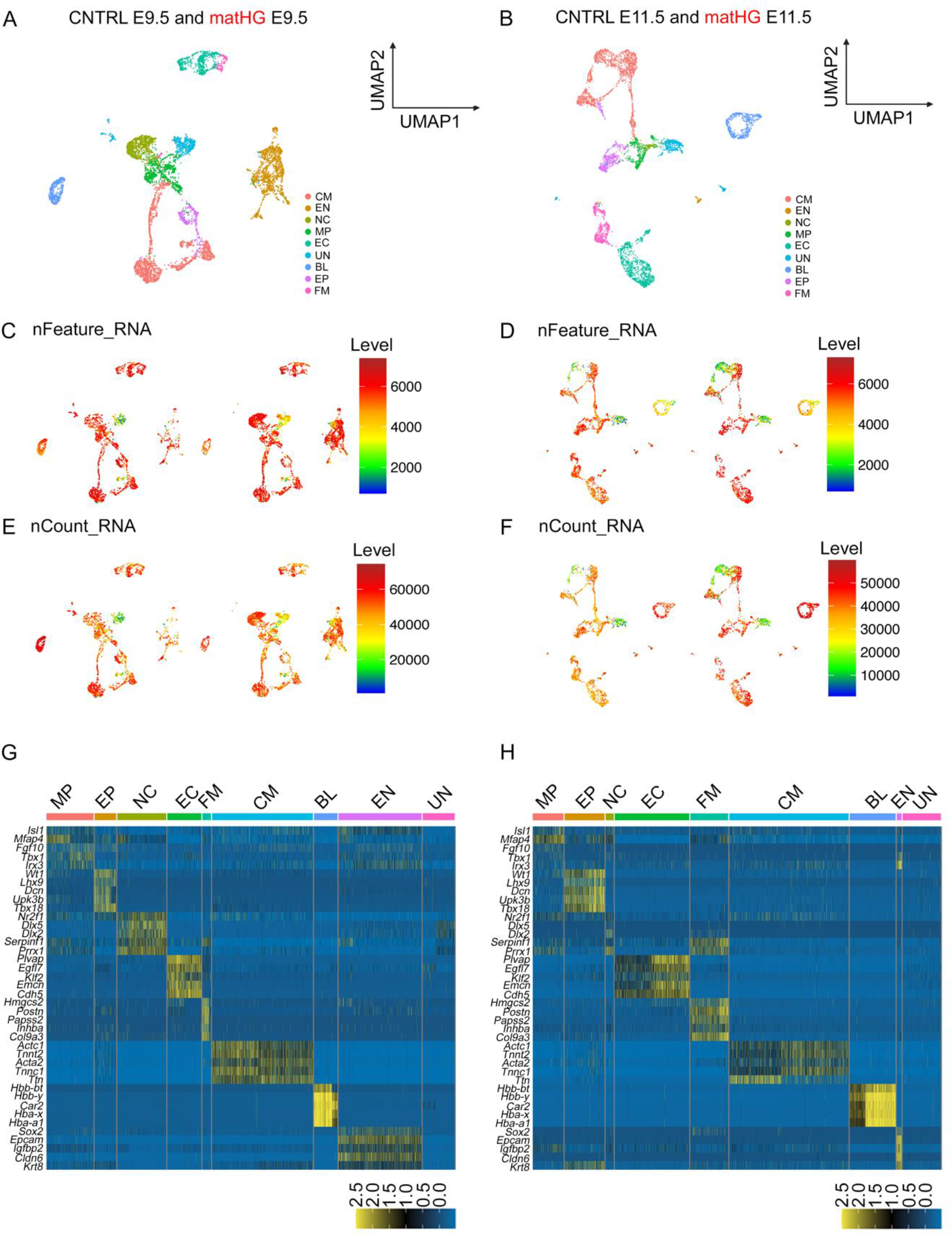
Developmental stage specific unsupervised clustering of control and matHG-exposed scRNA-seq data **(A, B)** UMAP plots show distribution of nine clusters (CM, EN, NC, MP, EC, UN, BL, EP, FM) separated based on developmental stages (E9.5 and E11.5), when exposed to maternal CNTRL and HG environment. Colors indicate cluster identities. **(C-F)** Individual UMAP plots show the relationship between the number of genes (nFeature- RNA) and UMIs (nCount-RNA) in E9.5 and E11.5 samples subjected to intrauterine CNTRL and matHG environment. **(G, H)** Unsupervised clustering shows nine clusters and the top five marker genes per cluster in the heatmap. Normalized log expression levels are shown in yellow (high expression) and dark blue (low expression). CM, cardiomyocytes, EN, endoderm, NC, neural crest, MP, multipotent progenitor, EC, endocardial/endothelial, UN, unknown, BL, blood, EP, epicardial, FM, fibromesenchymal, CNTRL, control, HG, hyperglycemia, UMI, unique molecular identifiers.

**Fig. S3.**
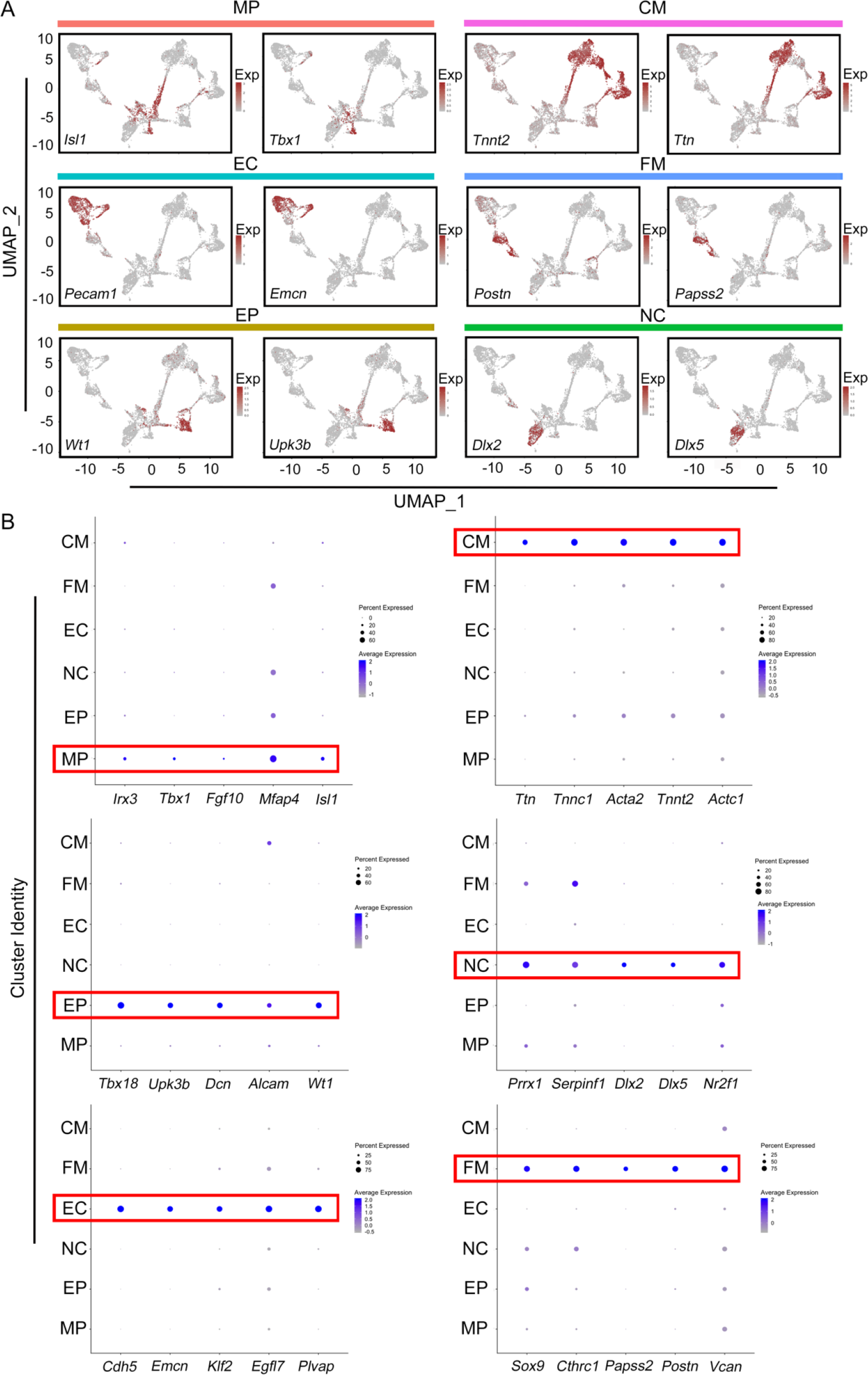
Cluster identification based on gene expression profile **(A)** UMAP plots illustrating cluster-specific expression of highly expressed marker genes applied to classify six broadly defined cardiac cell populations. The scale indicates Z-scored expression values (red = high expression, grey = no expression). **(B)** Dot plots showing the expression of known lineage marker genes across MP, EP, NC, EC, FM, and CM clusters. Each dot is sized to represent the percentage of cells of each type expressing the marker gene and colored to represent the average expression of each marker gene across all cells, as shown in the key (dark blue = high expression and light blue = low expression). MP, multipotent progenitor, EP, epicardial, NC, neural crest EC, endocardial/endothelial, FM, fibromesenchymal and CM, cardiomyocytes.

**Fig. S4.**
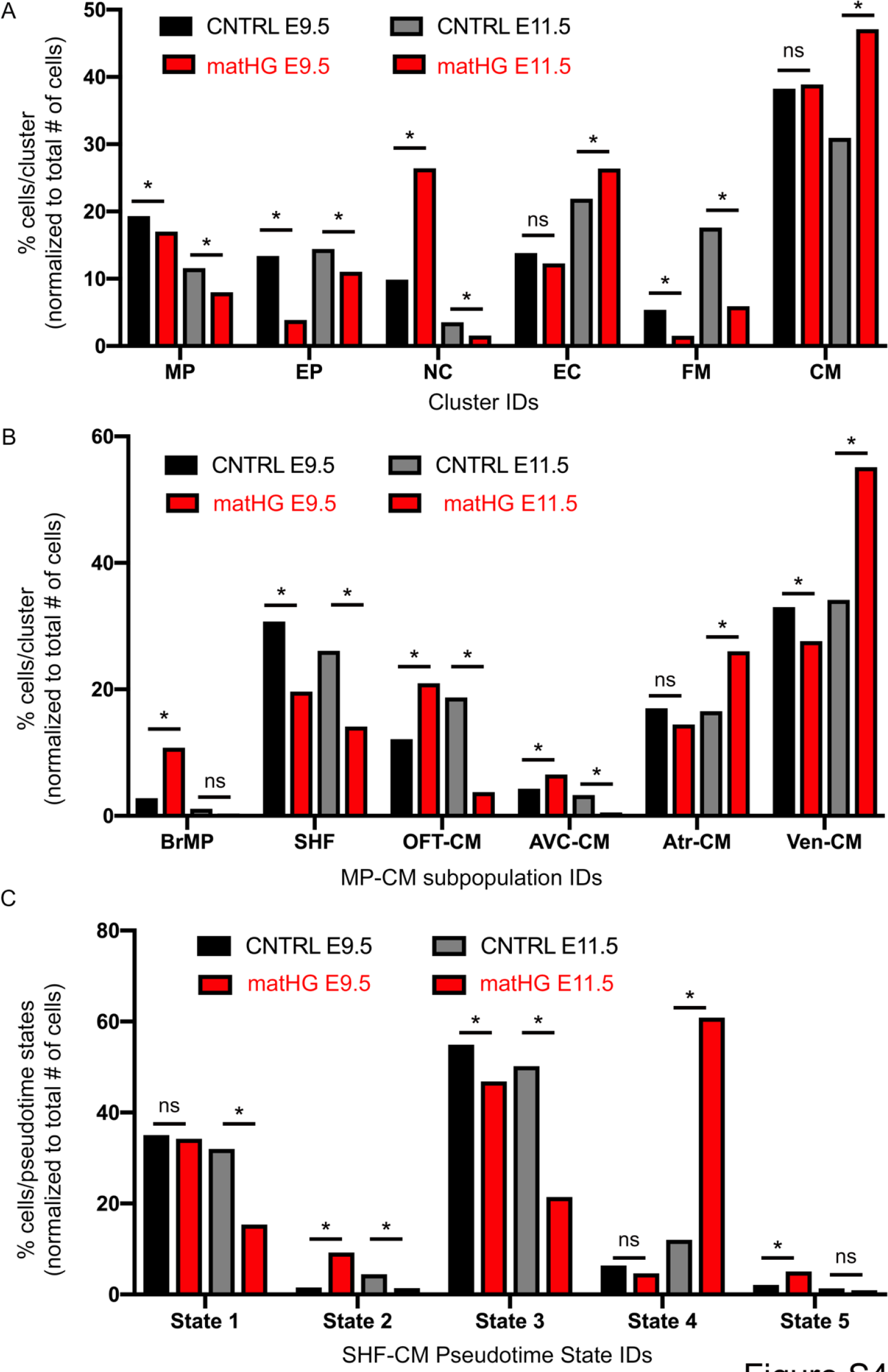
Maternal hyperglycemia alters the proportion of cells in each cluster and pseudotime states in developing embryonic hearts: **(A-C)** Graphs show the proportion of cell number in each cluster (A), MP-CM subclusters (B) and pseudotime states (C) from CNTRL and matHG exposed E9.5 and E11.5 hearts. For cell number comparisons, cells were normalized to the total number of cells analyzed. Statistical significance between groups determined by Fisher exact test. ns= non-significant and * indicates two-tailed p-value ≤ 0.05.

**Fig. S5.**
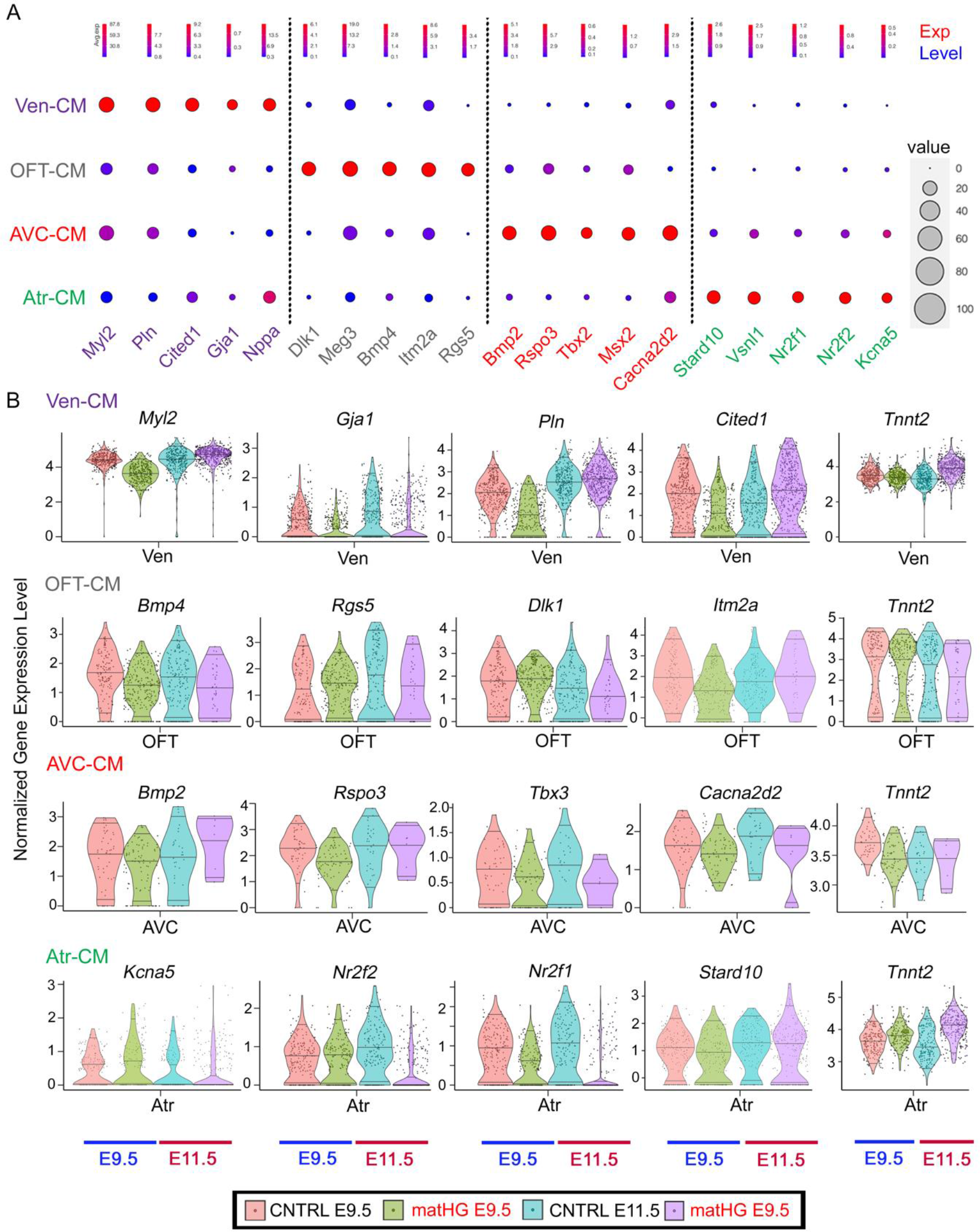
Subclustering cardiomyocyte subpopulations and effect of matHG on gene expression levels **(A)** Dot plots showing the expression of known marker genes across CM subtypes (Ven, OFT, AVC, and Atr-CMs). Each dot is sized to represent the proportion of cells of each type expressing the marker gene and colored to represent the average expression of each marker gene across all as shown in the key (red = high expression and dark blue = low expression). **(B)** Violin plots representing normalized gene-expression levels in each CM subclusters from CNTRL and matHG-exposed E9.5 and E11.5 hearts. Each dot in the violin plot represents individual cells. Ven, ventricular, OFT, outflow tract, AVC, atrioventricular canal, and Atr, atrial, CM, cardiomyocytes, CNTRL, control and HG, hyperglycemia.

**Fig. S6.**
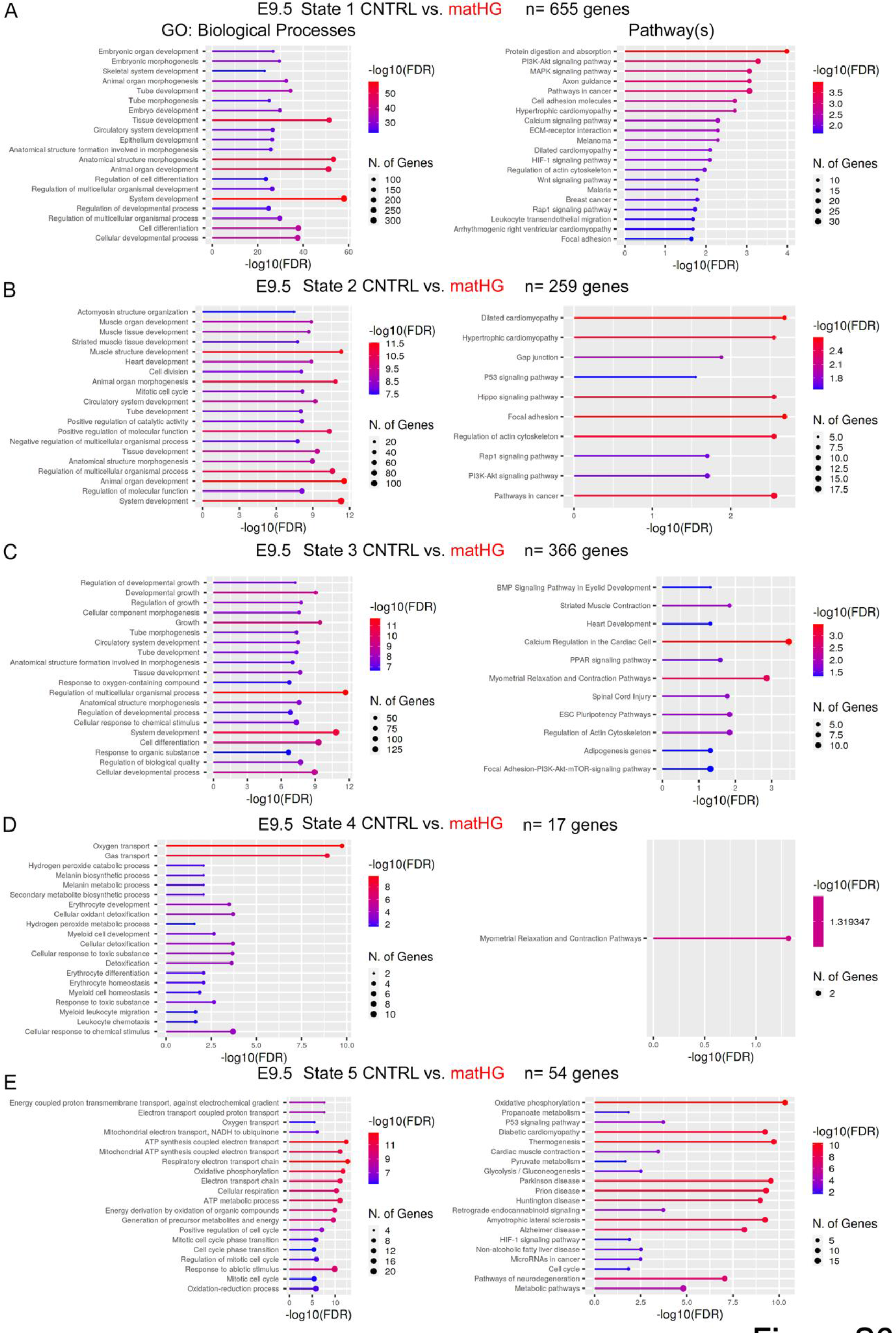
Transcriptomic changes in MP-CM subpopulations at E9.5 across the pseudotime trajectories **(A-E)** Lollipop charts illustrating the top 20 GO-terms for Biological Processes and Pathways, sorted as descending negative logarithmic adjusted p-value of enrichment analysis (-log10(FDR)) shown in the key. Numbers in circles represent the number of DEGs matched to a specific GO term. Five pseudotime states were compared between CNTRL and matHG exposed E9.5 MP-CM subclusters. GO, gene ontology, FDR, false discovery rate, DEGs, differentially expressed genes, CNTRL, control, HG, hyperglycemia.

**Fig. S7.**
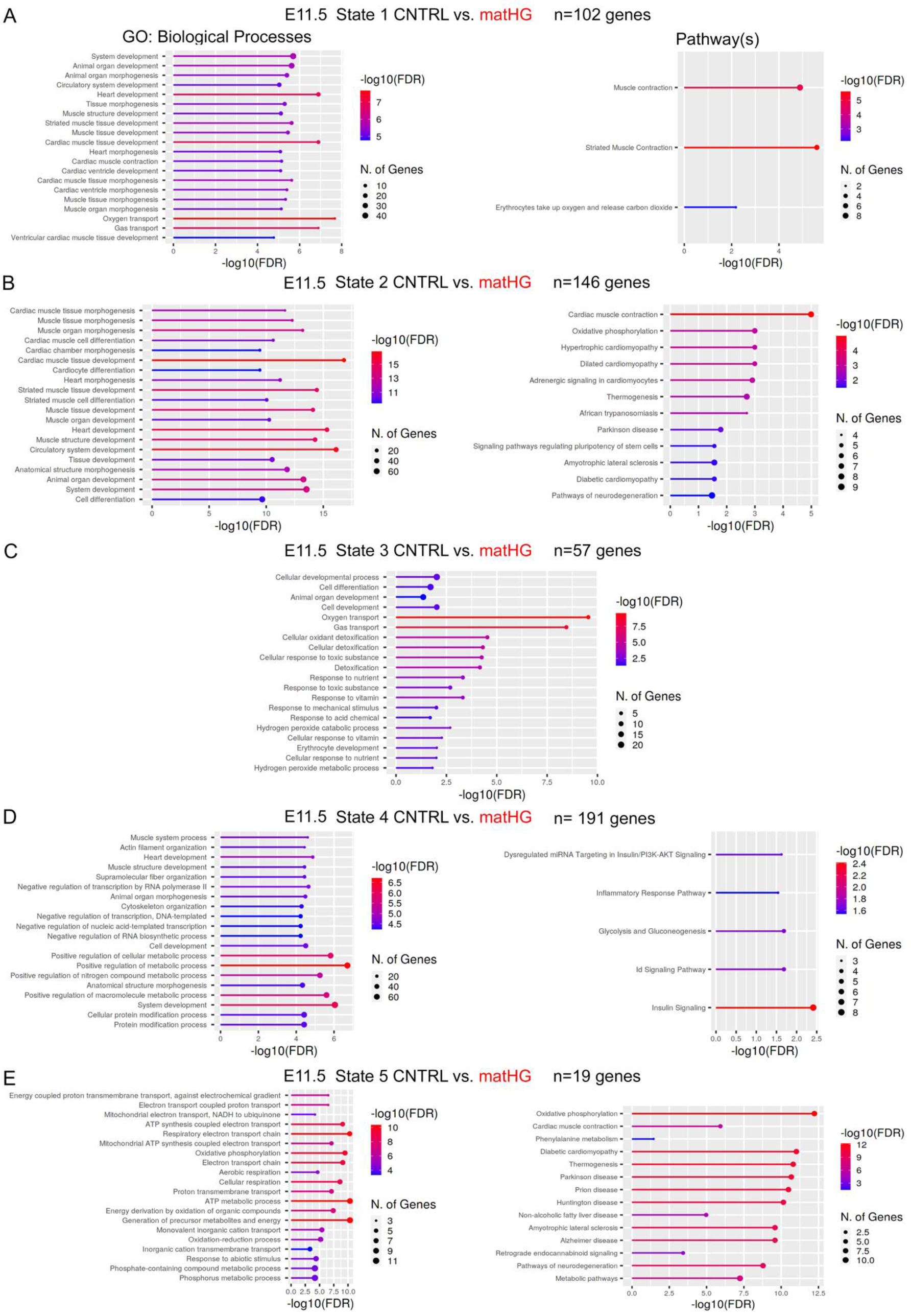
Transcriptomic changes in MP-CM subpopulations at E11.5 across the pseudotime trajectories **(A-E)** Lollipop plots representing the top 20 GO-terms for Biological Processes and Pathways, sorted as descending negative logarithmic adjusted p-value of enrichment analysis (-log10(FDR)) shown in the key. Numbers in circles represent the number of DEGs matched to a specific GO term. Five pseudotime states were compared between CNTRL and matHG exposed E11.5 MP-CM subclusters. No significantly enriched pathways were noted in cells present in State 3 at E11.5. GO, gene ontology, FDR, false discovery rate, DEGs, differentially expressed genes, CNTRL, control, HG, hyperglycemia.

**Fig. S8.**
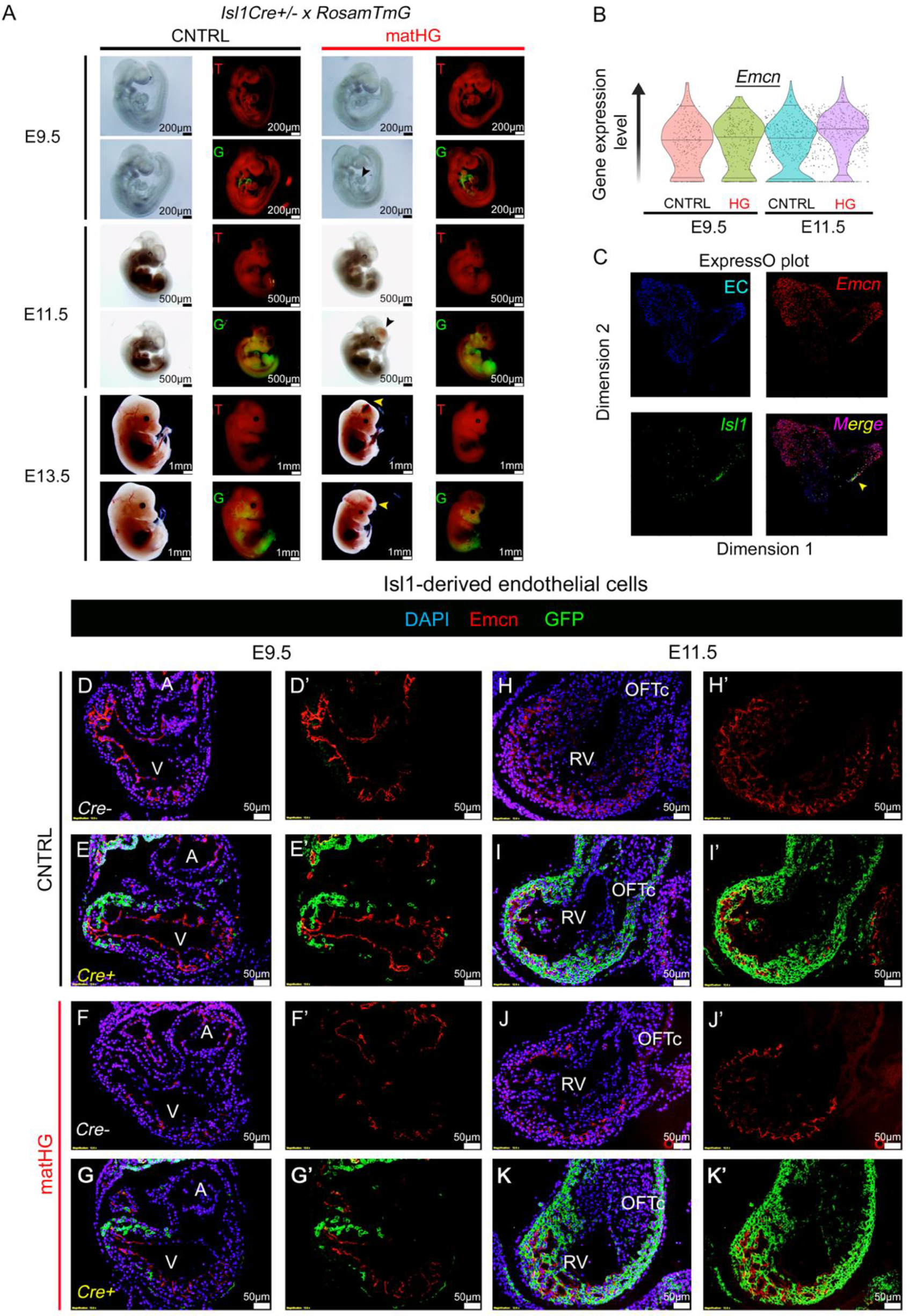
*Isl1*-cell fate mapping studies under maternal hyperglycemic exposure **(A)** Whole-mount brightfield and fluorescence images represent *Isl1Cre^-^; Rosa^mTmG/+^* (T; red, tdTomato^+^) and *Isl1Cre^+^; Rosa^mTmG/+^* (G; green, GFP^+^) littermate control hearts at E9.5, E11.5 and E13.5 stages exposed to CNTRL and matHG environment. GFP^+^ expression follows the endogenous pattern of *Isl1* expression and genetically label daughter cells derived from *Isl1*. MatHG-exposed *Isl1Cre^+^; Rosa^mTmG/+^* embryos result in intracerebral hemorrhage and exencephaly at E11.5 and E13.5 (shown in black and yellow arrowheads). **(B)** Violin plots show expression of known endothelial cell marker, *Emcn* expression in E9.5 and E11.5 scRNA-seq data. **(C)** ExpressO plot illustrating the presence of *Isl1^+^Emcn^+^* cells in EC cluster (yellow arrowheads). **(D-K)** Immunofluorescent images show GFP (green) and Emcn (red) protein expression in CNTRL and matHG-exposed Cre- and Cre+ littermate controls. Nuclei stained with DAPI shown in blue. **D’-K’** show *Isl1*-derived Emcn^+^GFP^+^ ECs (in yellow). CNTRL, control, HG, hyperglycemia, EC, endocardial/endothelial. Scale bars: **(A)** 200μm (E9.5), 500μm (E11.5) and 1mm (E13.5) and **(D-K)** 50μm.

**Fig. S9.**
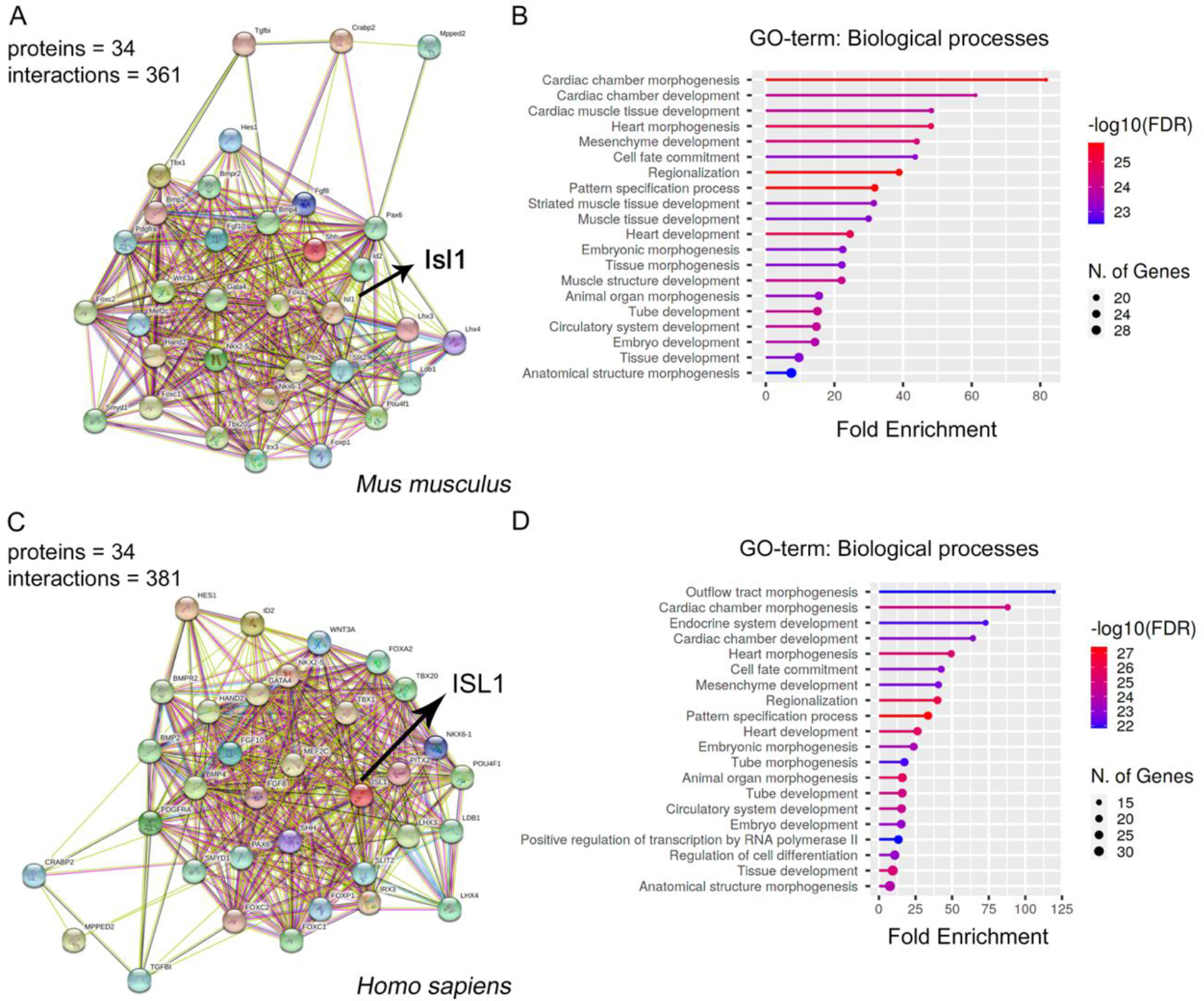
Isl1-mediated protein-protein interaction network **(A, C)** Isl1-GRN created in ShinyGOv0.741 with 34 proteins listed from STRINGdb and Brian Black, Semin Cell Dev Biol (2007)^24^ mapped for *Mus musculus* and *Homo sapiens*. STRING analysis of the protein interaction network showing predicted associations of the Isl1-interacting proteins. Line thickness indicates the strength of data support (STRING v11.5). Arrows indicate the location of Isl1 in the PPI network. **(B, D)** Species specific lollipop plots representing the top 10 GO-terms for Biological Processes sorted as descending negative logarithmic adjusted P-value of fold enrichment analysis shown in the key. Numbers in circles represent the number of genes matched to a specific GO term. PPI, protein-protein interaction, GO, gene ontology.

**Fig. S10.**
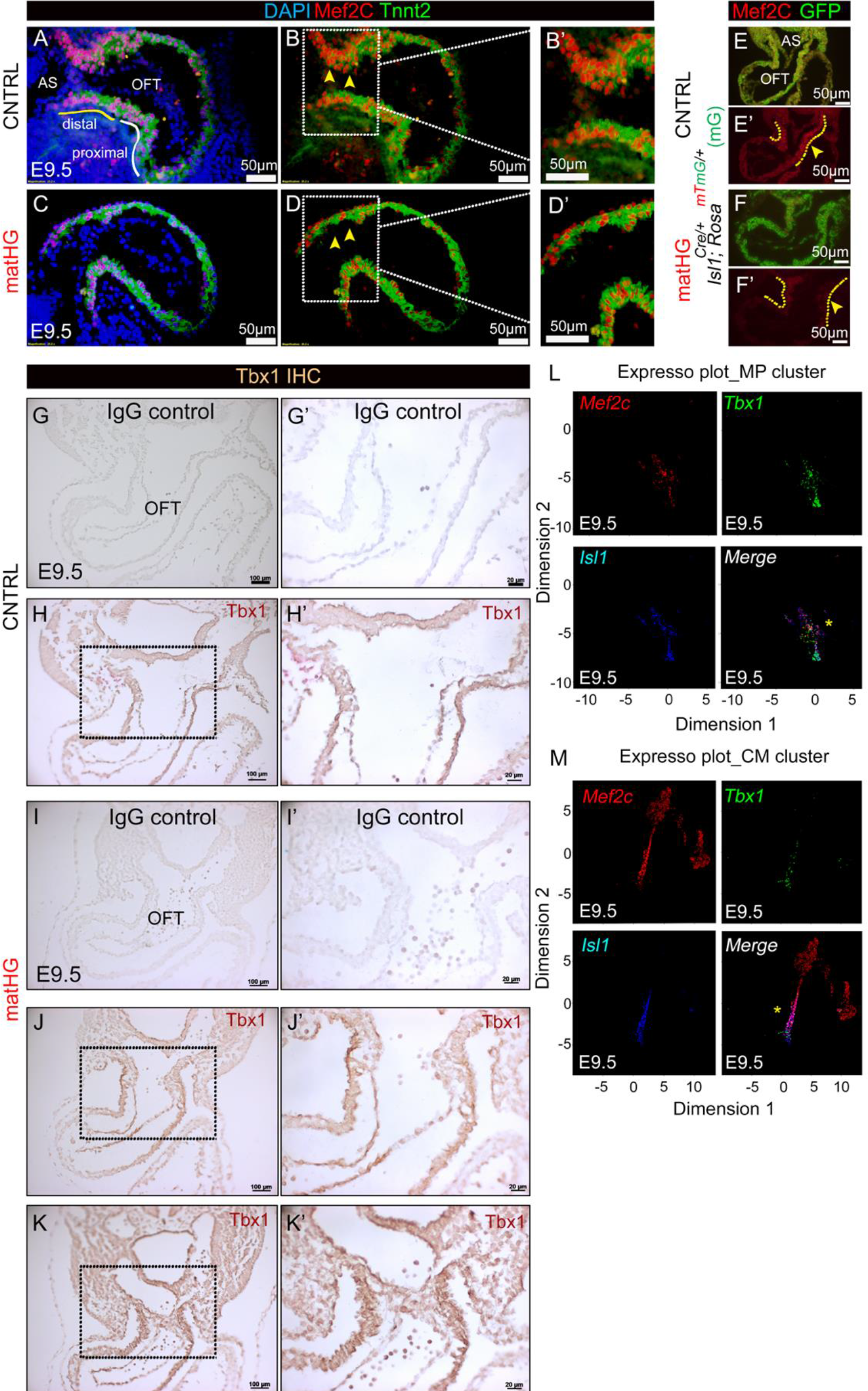
Expression of cardiac progenitor markers in response to maternal hyperglycemia **(A-D)** Immunofluorescent co-staining of Mef2c (red) and Tnnt2 (green) protein expression in CNTRL vs. high-matHG-exposed E9.5 *wt* hearts (in two independent embryos/group). Yellow and white solid lines in **(A)** indicate distal and proximal OFT. White dotted lines in **B** and **D** indicate high magnification images shown in **B’** and **D’** respectively. Nuclei stained with DAPI shown in blue. **(E, F)** Mef2c (red) and GFP (green) protein expression shown in E9.5 *Isl1Cre^+^; Rosa^mTmG/+^* (mG) hearts exposed to CNTRL and matHG environment (in three independent embryos/group). Yellow arrowheads and dotted lines indicate downregulation of Mef2c in the OFT. **(G-K)** Tbx1 IHC staining in CNTRL vs. matHG exposed E9.5 *wt* hearts (in three independent embryos/group). **G’-K’** show higher magnification images of **G-K** (indicated by black rectangular boxes). **G** and **I** display mock IgG (negative) control and **H, J** and **K** show upregulation of Tbx1 protein expression in the distal OFT. **(L, M)** ExpressO plots show transcript expression of *Mef2c, Tbx1,* and *Isl1* in MP and CM clusters. Yellow asterisks indicate merged expression of these progenitor cell markers in E9.5 scRNA-seq data. CNTRL, control, HG, hyperglycemia, wt, wildtype, OFT, outflow tract, IHC, immunohistochemistry. Scale bars: **A-F**: 50μm, **G-K**: 100μm and **G’-K’**: 50μm.

**Fig. S11.**
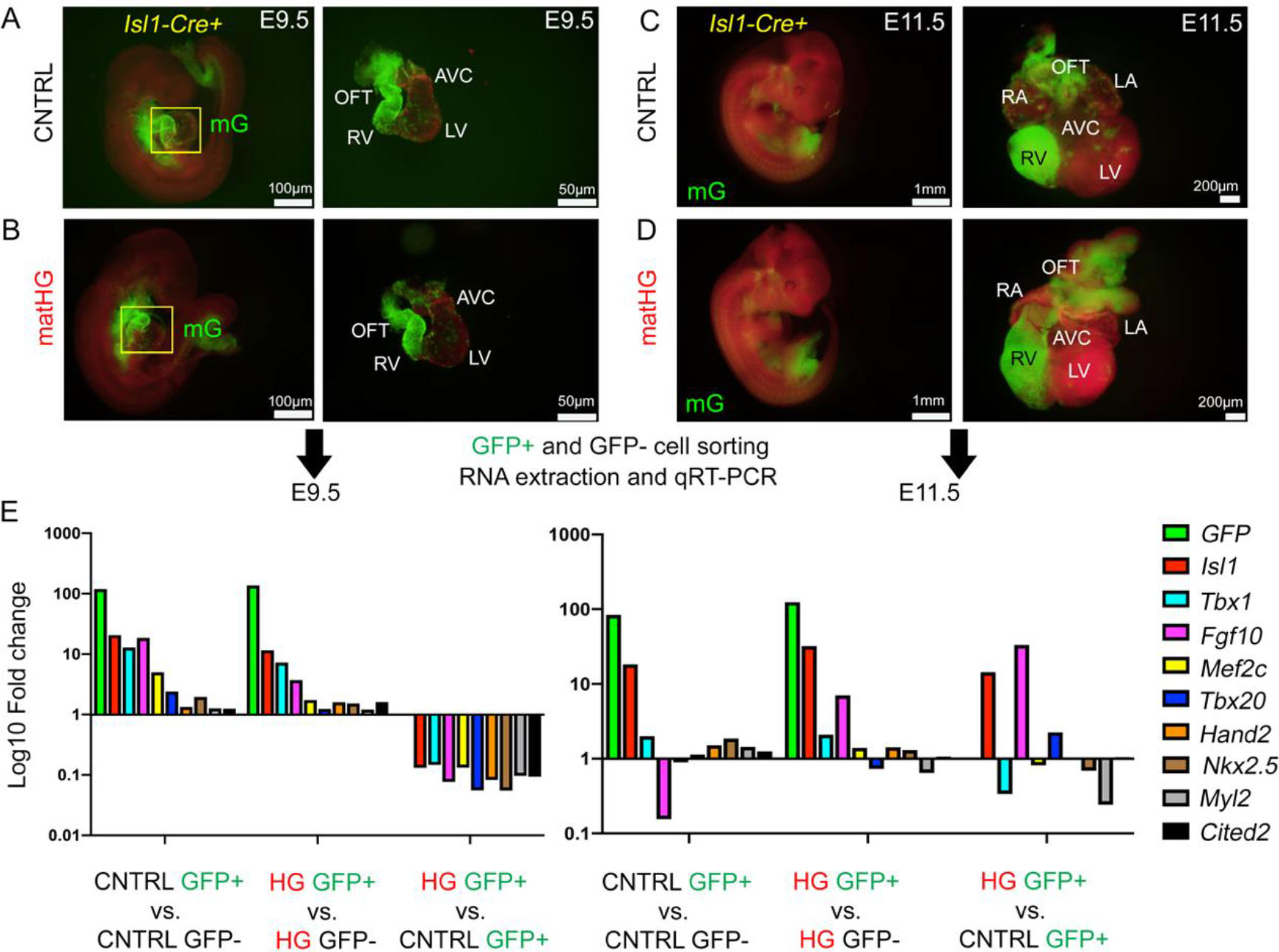
Expression of cardiac progenitor markers using flow sorted *Isl1*-derived cells in response to maternal hyperglycemia **(A-D)** Fluorescence microscopy images show the GFP expression in CNTRL and matHG exposed E9.5 and E11.5 *Isl1Cre^+^; Rosa^mTmG/+^* (mG) embryos and microdissected hearts (n≥3 embryos/timepoint). **(E, F)** Whole hearts were dissociated to single cells by enzymatic digestion. GFP^+^ and GFP^-^ cells sorted using FACS and relative gene expression levels (Log_10_ fold change) were measured using qRT-PCR analysis. Comparisons between CNTRL GFP^+^ vs. GFP^-^ and matHG GFP^+^ vs. GFP^-^ show enriched expression of GFP and other SHF/CM markers (*Isl1, Tbx1, Fgf10, Mef2c, Tbx20, Hand2, Nkx2.5, Myl2 and Cited2*) in FACS sorted GFP^+^ fraction under both maternal conditions. The comparisons of gene expression between CNTRL GFP^+^ and matHG GFP^+^ group demonstrate significant downregulation of SHF/CM markers at E9.5. Gene expression of cardiac progenitor markers *Isl1*, *Fgf10* and *Tbx20* were upregulated in matHG-exposed E11.5 GFP^+^ fraction. CNTRL, control, HG, hyperglycemia, FACS, Fluorescence-Activated Cell Sorting, qRT-PCR, quantitative real time polymerase chain reaction, SHF, second heart field, CM, cardiomyocytes.

**Fig. S12.**
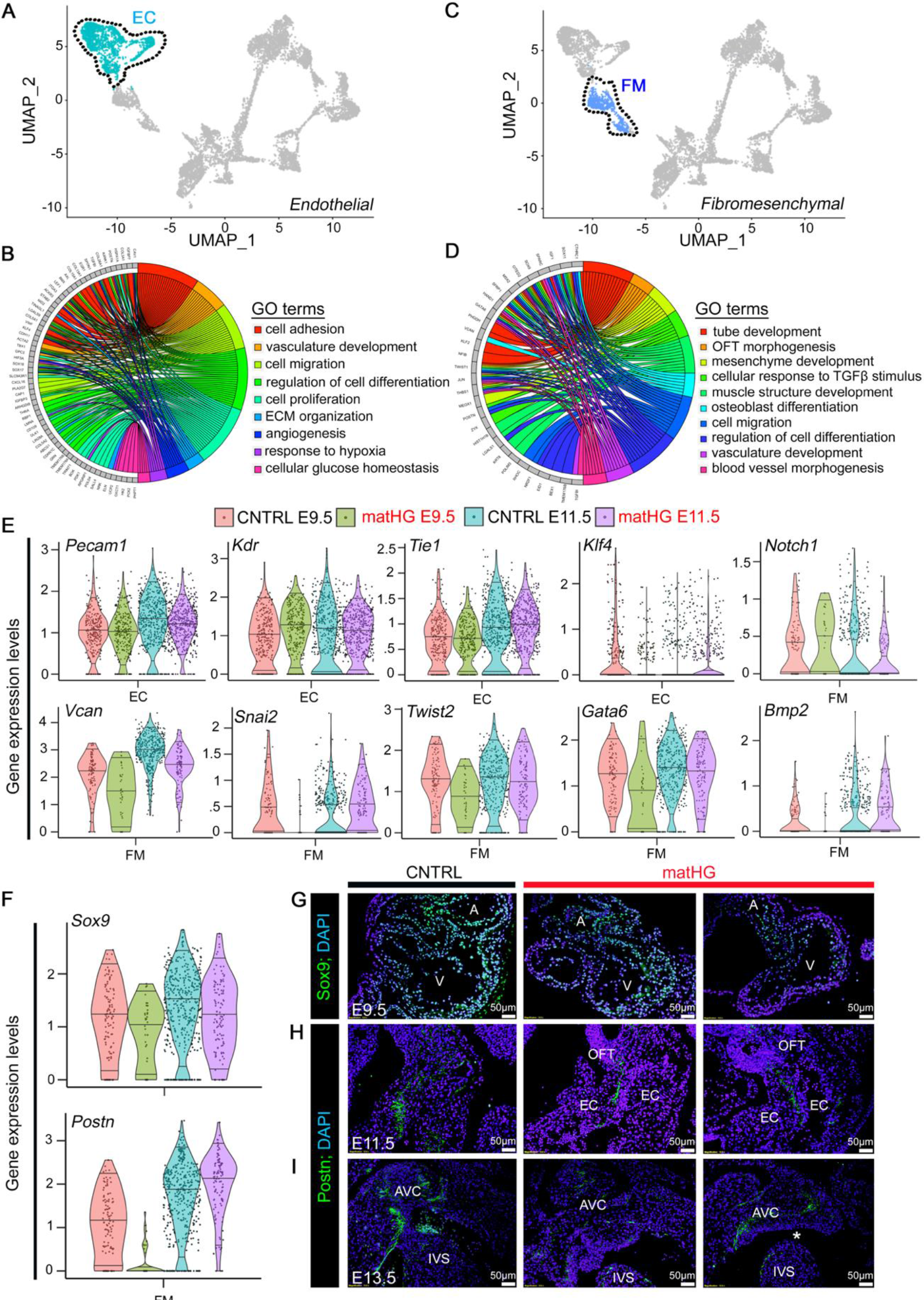
ScRNA-seq reveals transcriptional changes in endocardial/endothelial and mesenchymal cell populations exposed to maternal hyperglycemia **(A, B)** UMAP plots represent the EC and FM clusters from E9.5 and E11.5 embryonic hearts subjected to CNTRL and matHG. **(C, D)** GOplots represent the analysis of the GO terms enriched among the DEGs in E9.5 and E11.5 EC and FM clusters (DEG cutoff: Log2Foldchange ≥ 1 or ≤ -1 and P_adjusted_ ≤ 0.05). The left side of the circle displays the gene, and the right side shows the GO-term associated biological processes. The assorted colors represent different GO terms. **(E)** Violin plots show the normalized expression levels of highly variable genes in EC (*Pecam1, Kdr, Tie1, and Klf4*) and FM (*Vcan, Snai2, Twist2, Gata6, Bmp2, and Notch1*) clusters from CNTRL and matHG-exposed E9.5 and E11.5 embryonic hearts. **(F)** Violin plots illustrating *Sox9* and *Postn* gene-expression in E9.5 and E11.5 FM populations exposed to CNTRL and matHG environment. **(G-I)** Panels of immunofluorescent images show protein expression of Sox9 (green) and Postn (green) at E9.5, E11.5 and E13.5 hearts exposed to CNTRL and matHG. Nuclei stained with DAPI shown in blue. White asterisk denotes presence of VSD at matHG-exposed E13.5 embryo. Nuclei stained with DAPI in blue. CNTRL, control, HG, hyperglycemia, GO, gene ontology, FM, fibromesenchymal, A, atria; V, ventricle; OFT, outflow tract; EC, endocardial/endothelial cushion; AVC, atrioventricular canal; IVS, interventricular septum, VSD, ventricular septal defect. Scale bars: **G-I**: 50μm.

**Fig. S13.**
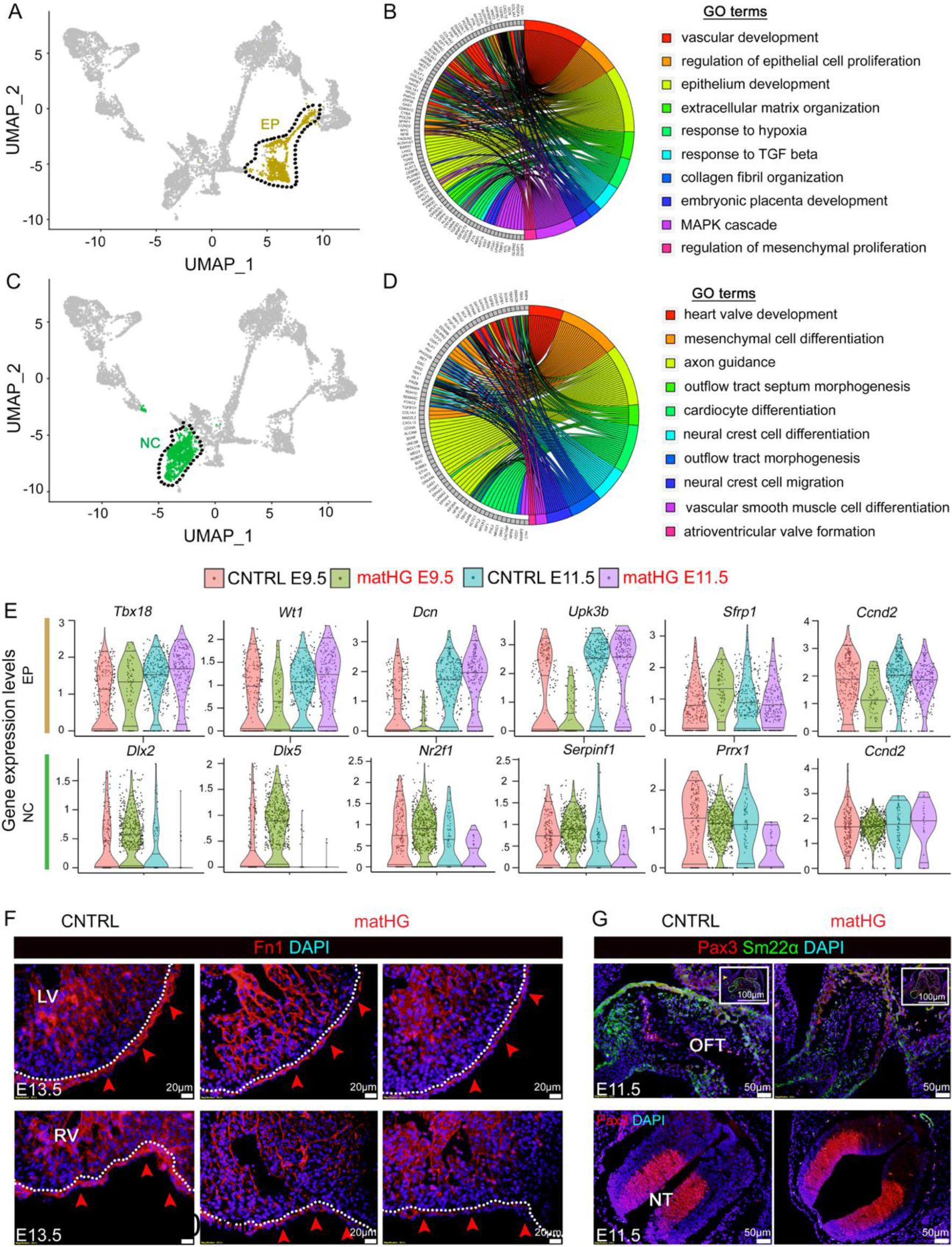
ScRNA-seq reveals transcriptional changes in epicardial and cardiac neural crest cell populations exposed to maternal hyperglycemia **(A, B)** UMAP plots represent the EP and NC clusters from E9.5 and E11.5 embryonic hearts subjected to CNTRL and matHG. **(C, D)** GOplots represent the GO terms enriched among the DEGs in E9.5 and E11.5 EP and NC clusters (DEG cutoff: Log2Foldchange ≥ 1 or ≤ -1 and P_adjusted_ ≤ 0.05). The left-side of the circle displays the gene, and the right-side shows GO-term associated biological processes. The assorted colors represent GO terms. **(E)** Violin plots show the normalized expression levels of highly variable genes in EP (*Tbx18, Wt1, Dcn, Upk3b, Sfrp1, and Ccnd2*) and NC (*Dlx2, Dlx5, Nr2f1, Serpinf1, Prrx1, and Ccnd2*) clusters from CNTRL and matHG-exposed E9.5 and E11.5 embryonic hearts. **(F)** Panels of immunofluorescent images show protein expression of *Fn1* (red arrowheads) in the EP from low to high-matHG exposed E13.5 LV and RV compared to CNTRL (indicated by dotted white line). **(G)** Panels of immunofluorescent images illustrate the protein expression of NC-markers, Pax3 (red) and Sm22α (green) in E11.5 OFT exposed to matHG vs. CNTRL. Insets (white square boxes) show positive Pax3 expression in the NT. Nuclei stained with DAPI in blue. CNTRL, control, HG, hyperglycemia, GO, gene ontology, EP, epicardial, NC, neural crest, LV, left ventricle, RV, right ventricle, OFT, outflow tract, NT, neural tube. Scale bars: **F**: 20μm and **G**:50μm.

## Supplemental Tables

Table S1. List of DEGs in CNTRL vs. matHG-exposed E9.5 hearts.

Table S2. List of GO terms associated with DEGs in CNTRL vs. matHG-exposed E9.5 hearts

Table S3. List of DEGs in CNTRL vs. matHG-exposed E11.5 hearts

Table S4. List of GO terms associated with DEGs in CNTRL vs. matHG-exposed E11.5 hearts.

Table S5. List of DEGs in CNTRL vs. matHG exposed E9.5 MP-CM subclusters

Table S6. List of DEGs in CNTRL vs. matHG-exposed E11.5 MP-CM subclusters

Table S7. List of GO-terms in E9.5 and E11.5 MP-CM subclusters

Table S8. List of DEG analysis in CNTRL vs. matHG exposed E9.5 MP-CM pseudotime states

Table S9. List of DEG analysis in CNTRL vs. matHG exposed E11.5 MP-CM pseudotime states

**Table S10:**
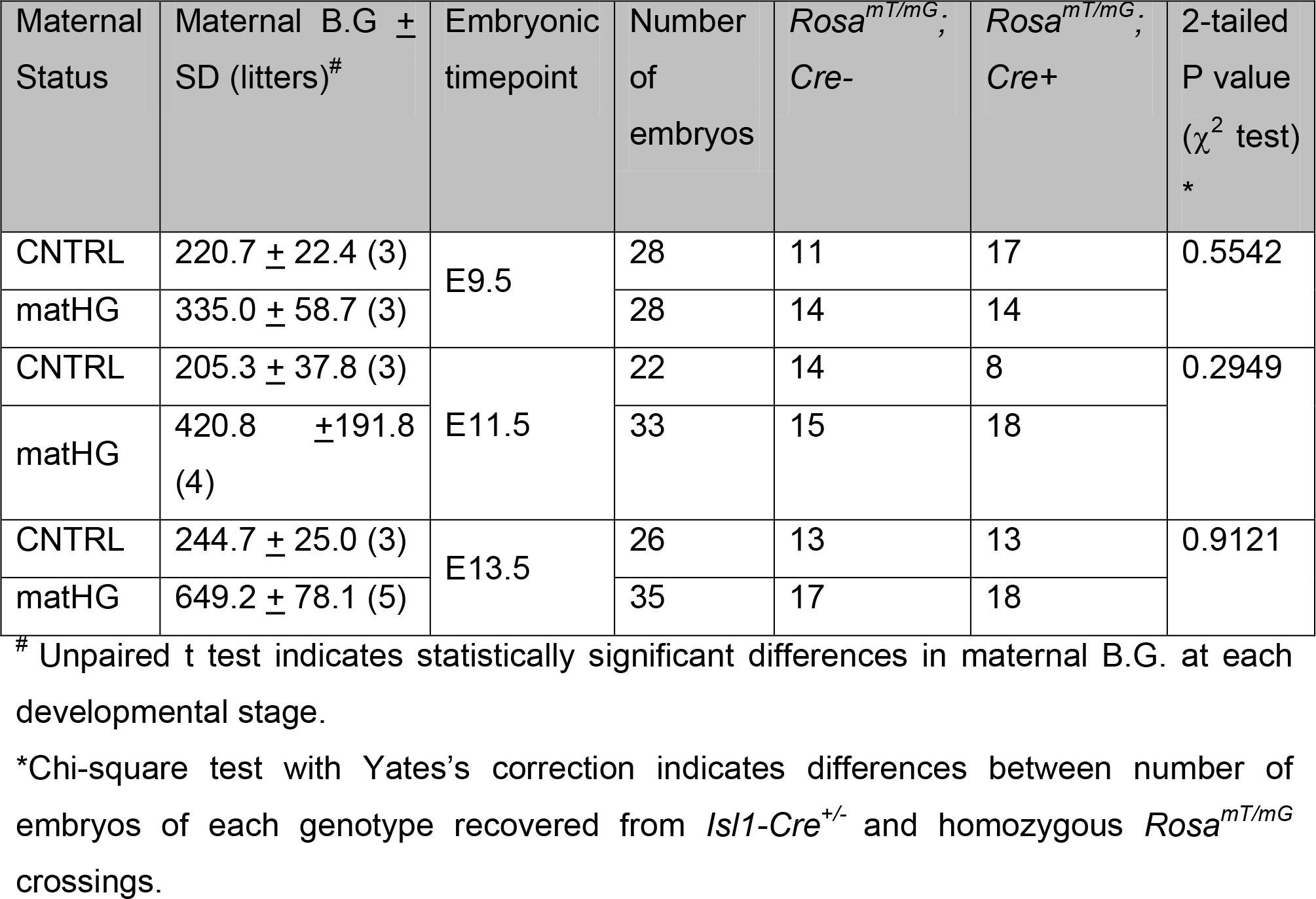
Distribution of Isl1-Cre^+/-^; Rosa^mT/mG^ embryos in the setting of control and matHG environment

**Table S11.**
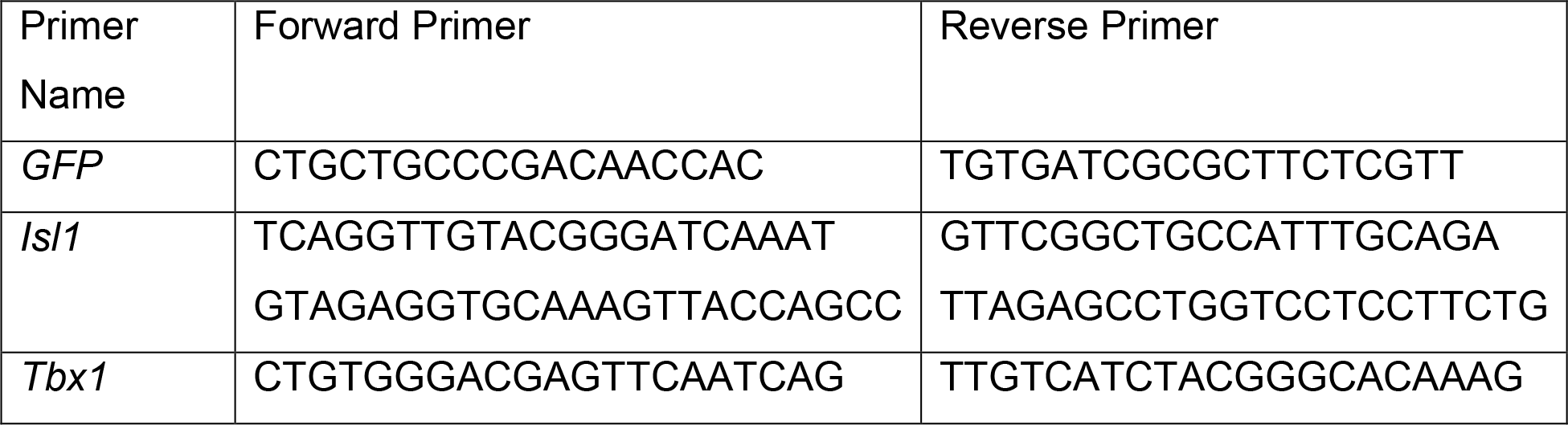
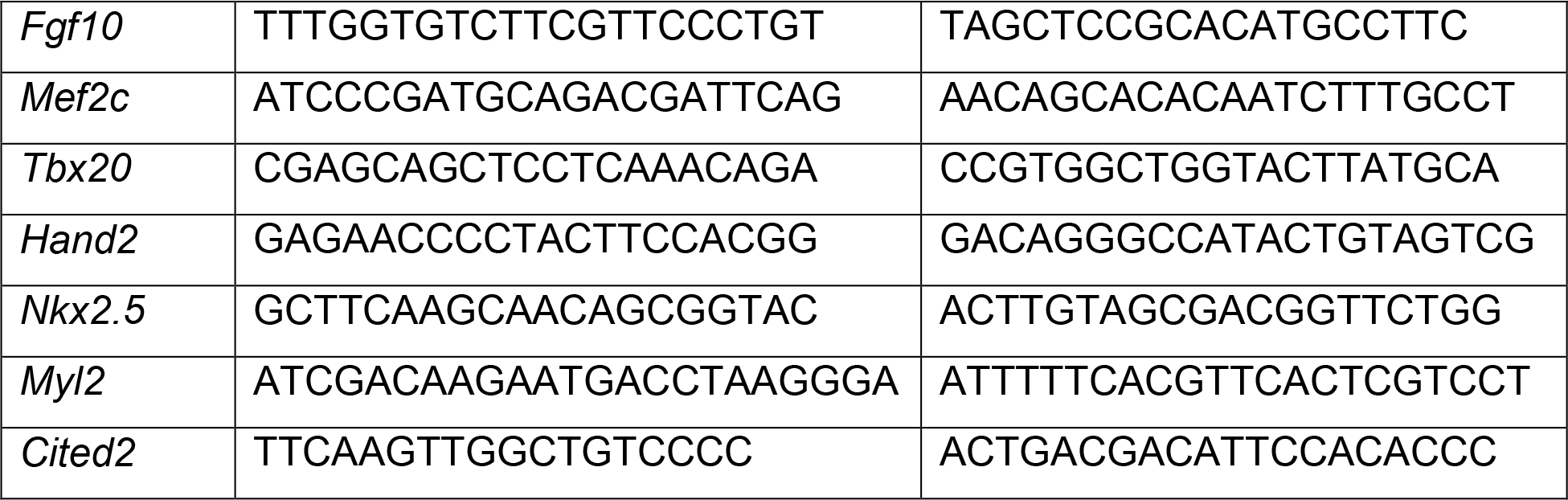
Oligonucleotide sequences

## Notes

### Competing Interest Statement

The authors have declared no competing interest.

### Summary of Updates

New data has been added to the revised manuscript. The current changes does not change the final inference of the paper.

